# Resting mitochondrial complex I from *Drosophila melanogaster* adopts a helix-locked state

**DOI:** 10.1101/2022.11.01.514701

**Authors:** Abhilash Padavannil, Anjaneyulu Murari, Shauna-Kay Rhooms, Edward Owusu-Ansah, James A. Letts

## Abstract

Respiratory complex I is a proton-pumping oxidoreductase key to bioenergetic metabolism. Biochemical studies have found a divide in the behavior of complex I in metazoans that aligns with the evolutionary split between Protostomia and Deuterostomia. Complex I from Deuterostomia including mammals can adopt an off-pathway “deactive” state, whereas complex I from Protostomia cannot. The presence of off-pathway states complicates the interpretation of structural results and has led to considerable mechanistic debate. Here we report the structure of mitochondrial complex I from the thoracic muscles of the model protostomian *Drosophila melanogaster*. We show that, although *D. melanogaster* complex I (*Dm*-CI) does not deactivate the resting state of *Dm*-CI adopts multiple conformations. We identify a new helix-locked open state in which an N-terminal α-helix on the NDUFS4 subunit wedges between the peripheral and membrane arms. Comparison of the *Dm*-CI structure and conformational states to those observed in bacteria, yeast and mammals provides insight into the roles of subunits across organisms, explains why *Dm*-CI does not deactivate and reveals incompatibilities with current mechanistic models of complex I turnover. Additionally, the *Dm*-CI structure and novel regulatory mechanism will allow for the development of more selective pesticides for agriculture and human disease.

## Introduction

The final stage of eukaryotic cellular respiration occurs in the mitochondria. A series of large membrane protein complexes that reside in the inner mitochondrial membrane (IMM) form the oxidative phosphorylation (OXPHOS) electron transport chain (ETC) that catalyzes the terminal respiratory reactions. ETC complexes are a series of redox-coupled H^+^ pumps that connect oxygen consumption to adenosine triphosphate (ATP) synthesis by the generation of a proton motive force (pmf) across the IMM that is used to power the ATP synthase complex. In metazoans, the generation of the pmf is driven by the transfer of electrons from reduced substrates (NADH and succinate) to O_2_ via four ETC complexes (Complexes I-IV, CI-IV) and electron-carriers ubiquinone (coenzyme Q, CoQ) and cytochrome *c* (cyt *c*). In addition to their independent existence, ETC complexes can form higher order structures known as supercomplexes (SCs) (Letts and Sazanov, 2017; Schägger and Pfeiffer, 2000). Across species, the most common SCs are formed between CI, a dimer of CIII (CIII_2_) and CIV (SC I+III_2_+IV) (Letts et al., 2016); CI and CIII_2_ alone (SC I+III_2_) (Letts et al., 2019); and CIII_2_ and CIV (SC III_2_+IV) (Hartley et al., 2018; Rathore et al., 2018; Vercellino and Sazanov, 2021). The functional advantage of SC formation remains undefined, but they are found across eukaryotes often as the most abundant form of the ETC complexes (Davies et al., 2018; Schägger and Pfeiffer, 2001; Zhou et al., 2022).

CI couples the transfer of electrons from NADH to CoQ to the pumping of four H^+^ across the IMM (Galkin et al., 1999; Jones et al., 2017). It has an “L” shaped structure consisting of two arms: a peripheral arm (PA) that extends into the matrix and a membrane arm (MA) that is embedded in the IMM. CI accepts electrons from NADH onto an FMN co-factor near the distal tip of the PA and transfers them to CoQ via a series of seven iron-sulfur (FeS) clusters. With few exceptions, the catalytic core of 14 subunits is conserved across species and contain all the redox cofactors and active sites required for catalysis (Hirst, 2013; Sazanov, 2015). In addition to the core subunits, eukaryotic CI has varying numbers of accessory subunits, e.g., 29 in the yeast *Yarrowia lipolytica* and 31 in mammals (Carroll et al., 2003; Letts and Sazanov, 2015; Padavannil et al., 2022; Parey et al., 2019). The accessory subunits are needed for assembly and stability of the complex (Garcia et al., 2017; Stroud et al., 2016) and in some cases may play an active role in regulating turnover (Padavannil et al., 2022).

The molecular mechanism of the coupling between the electron transfer and proton pumping has been the target of much research and debate (Chung et al., 2022a; Kampjut and Sazanov, 2022). Nonetheless, thanks to a plethora of high-resolution structures, a framework for the coupling mechanism is emerging in which specific conformational changes in CoQ binding site loops at the interface of the PA and MA initiate a wave of conformational changes upon CoQ reduction that propagate along the MA via a hydrophilic axis of amino acid residues resulting in H^+^ pumping (Kampjut and Sazanov, 2020; Kravchuk et al., 2022). Although this model is consistent with most mutagenesis and structural data, more experiments are needed to confirm the predictions of the model and examine possible variations in coupling and regulation across organisms (Klusch et al., 2021; Maldonado et al., 2020; Zhou et al., 2022).

Complicating the structural elucidation of the coupling mechanism is the fact that resting CI from opisthokonts (yeast and mammals) have been shown to exist in two distinct biochemical states known as the catalytically competent active (A) state and the off-pathway deactive (D) state (Gavrikova and Vinogradov, 1999; Maklashina et al., 2003). In yeast and mammals, CI spontaneously undergoes an A-to-D transition when exposed to physiological temperatures in the absence of reduced substrates (Babot et al., 2014). Biochemically, the D state is characterized by a solvent-exposed cysteine residue on the ND3 core subunit that can be modified by thiol-reactive agents such as N-ethyl maleimide (NEM) (Galkin et al., 2008). Modification of the cysteine residue traps CI in the D state. The presence of the D state is thought to protect cells from ROS-mediated damage due to CI reverse electron transport (Chouchani et al., 2013). It remains unclear whether conformations of CI in the presence (Kampjut and Sazanov, 2020) or absence (Agip et al., 2018; Blaza et al., 2018) of added substrates, which differ in the structure of ND3’s cysteine-containing loop and in the angle between the PA and MA, correspond to the A and D states of CI or if they are part of the CI’s catalytic cycle (Chung et al., 2022a; Kampjut and Sazanov, 2022).

Nonetheless, the A/D transition is not universally conserved across species (Maklashina et al., 2003). Biochemical characterization of CI from bacteria, protostomians and more recently the ciliate *Tetrahymena thermophila* have failed to detect a deactive state using the standard NEM approach (Maklashina et al., 2003; Zhou et al., 2022). Structural analysis of the *T. thermophila* CI demonstrated that its PA/MA interface is more extensive than in other species (Zhou et al., 2022), likely precluding the opening of the complex in the same manner observed in mammals (Agip et al., 2018; Kampjut and Sazanov, 2020) and *E. coli* (Kolata and Efremov, 2021; Kravchuk et al., 2022).

Further structural and biochemical analyses of CI in organisms without an A-to-D transition, e.g., insects or other protostomians, are needed to determine whether the observed conformational changes are catalytic or part of the off-pathway deactive state. Additionally, structures of CIs from organisms with differences in accessory subunits will reveal subunit function and novel ways in which CI can be regulated. Furthermore, CI is an established target for insecticides and agricultural pesticides (Murai and Miyoshi, 2016). Particularly with the emergence of *Drosophila suzukii* as a major pest for soft summer fruits (Tait et al., 2021), it is important to elucidate the structure of CI from a *Drosophila* species to identify unique aspects of CI in this genus. This structure will also inform on CI among insects more generally, including human disease vectors, which could be exploited for insecticide development.

*D. melanogaster* is a genetically tractable system, has a similar ETC to humans and is an emerging model for CI assembly and regulation (Garcia et al., 2017; Murari et al., 2022, 2021, 2020; Rhooms et al., 2019; Xu et al., 2019). *D. melanogaster* diverged from mammals (deuterostomians) after the evolution of bilaterians ∼600 million years ago (Figure 1A) (Dunn et al., 2014). Importantly, each protostomian CI that has been biochemically characterized lacks an A-to-D transition, whereas all fungal and deuterostomian CI characterized shows a clear A-to-D transition (Figure 1A) (Maklashina et al., 2003). Given that fungi diverged from metazoans ∼1300 million years ago, this indicates either the loss of an ancestral A-to-D transition in Protostomia or the evolution of distinct, though biochemically similar, A-to-D transitions in Fungi and Deuterostomia. Furthermore, *D. melanogaster* thoracic muscle mitochondria contain fewer supercomplexes than mammalian mitochondria (Garcia et al., 2017), indicating differences in the higher-order assembly of the ETC. Thus, to understand the diversity of CI biochemistry and assembly across opisthokonts, characterization of *D. melanogaster* CI (*Dm*-CI) is needed to compare with the available structures from fungi and mammals. Here we report the structure of resting CI from the thoracic muscles of the model protostomian *Drosophila melanogaster*.

**Figure 1.**
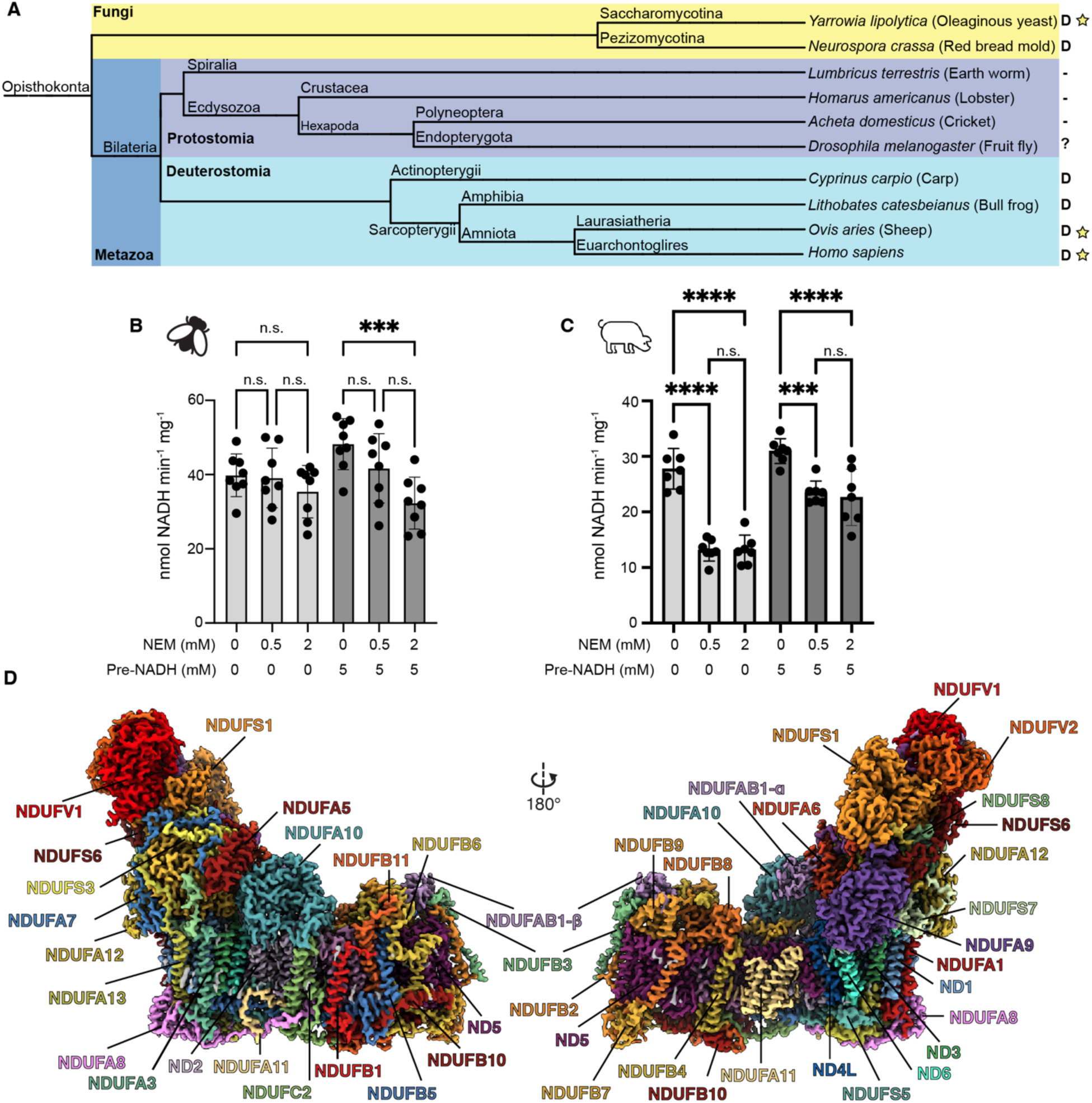
Evolution, biochemical characterization, and structure of *D. melanogaster* mitochondrial CI. (A) Dendrogram showing biochemically characterized CI from Opisthokonts. Distinct groups are highlighted with Fungi in yellow and metazoans in blue (Protostomia in dark blue and Deuterostomia in cyan). ‘D’ indicates the presence of biochemically characterized deactive state in the CI of the species. A minus sign indicates the absence of biochemically characterized deactive state in the CI of the species. A question mark indicates that the species has not been biochemically characterized for the presence of the deactive state. A star indicates that structures of CI from that species is currently available. (B C) Functional characterization of A-to-D transition in isolated mitochondrial membranes of *D. melanogaster* (B) and *S. scrofa* (C) by spectroscopic measurement of NADH dehydrogenase activity at 340 nm in the presence of the indicated concentrations of N-ethylmaleimide (NEM) with preincubation with 5 mM NADH or water. Individual values, average and SEM are shown, n = 7-8. Statistical analysis with ANOVA with Šídák’s multiple comparisons test. **, *p* < 0.01; ****, *p* < 0.0001, n.s. not significant. (D) Cryo-EM reconstruction of *Dm*-CI colored by subunit.

## Results

### *D. melanogaster* CI does not have an A-to-D transition

We assayed for the presence of an A-to-D transition in *Dm*-CI using the established NEM sensitivity assay on isolated *D. melanogaster* mitochondrial membranes (Figure 1B) (Babot et al., 2014; Galkin et al., 2008). In this assay, incubation of the mitochondrial membranes at 37°C in the absence of substrate should deactivate CI resulting in increased sensitivity to NEM inhibition. The impact of NEM incubation can be reduced by re-activation of the complex with a small amount of NADH prior to the addition of the NEM. When compared to mammalian mitochondrial membranes (*Sus scrofa*, Figure 1C), the *D. melanogaster* NADH oxidation rate is unaffected by NEM after incubation at elevated temperatures (Figure 1B and C). This is consistent with the lack of an A-to-D transition in *Dm*-CI as has been seen in the other protostomians, namely *Lumbricus terrestris* (earth worm), *Homarus americanus* (lobster) and *Acheta domesticus* (cricket) (Figure 1A) (Maklashina et al., 2003). Moreover, although the addition of NADH prior to NEM rescues activity in *Sus scrofa* (Figure 1C), pre-incubation with NADH sensitizes *Dm*-CI to NEM treatment, significantly reducing the rate of NADH oxidation after addition of 2 mM NEM (Figure 1B). This result is consistent with a model in which the ND3 cysteine remains buried in resting *Dm*-CI but becomes accessible during turnover (Kampjut and Sazanov, 2022). To better understand the functional differences between protostomian and deuterostomian CI, we solved the structure of *Dm*-CI by single particle cryogenic electron microscopy (cryoEM).

### Overall structure of mitochondrial CI from *D. melanogaster* thoracic muscle

*Dm*-CI was extracted from washed mitochondrial membranes using the mild detergent digitonin followed by exchange into amphipathic polymer A8-35 and enrichment by sucrose gradient ultracentrifugation (Figure 1-figure supplement 1A). Fractions containing CI activity, as assessed by blue-native polyacrylamide gel electrophoresis (BN-PAGE) in-gel activity (Figure 1-figure supplement 1B), were pooled and concentrated. These samples were applied directly to EM grids and used for cryoEM data collection (Table 1). The structure of *Dm*-CI was resolved to a nominal resolution of 3.3 Å (Figure 1D, Video 1, Figure 1-figure supplement 2, Table 1). This partially purified sample also contained particles of *Dm*-CIII_2_ and *Dm*-CV, albeit in insufficient numbers for high-resolution reconstruction (Figure 1-figure supplement 2). Consistent with previous studies on *Dm*-CI assembly (Garcia et al., 2017), but in contrast to what is seen in mammalian cardiac mitochondria of mammals, we did not observe significant amounts of SC I+III_2_ either biochemically on the BN-PAGE gels or as particles in the cryoEM data set (Figure 1-figure supplement 1B and 2).

**Figure 2.**
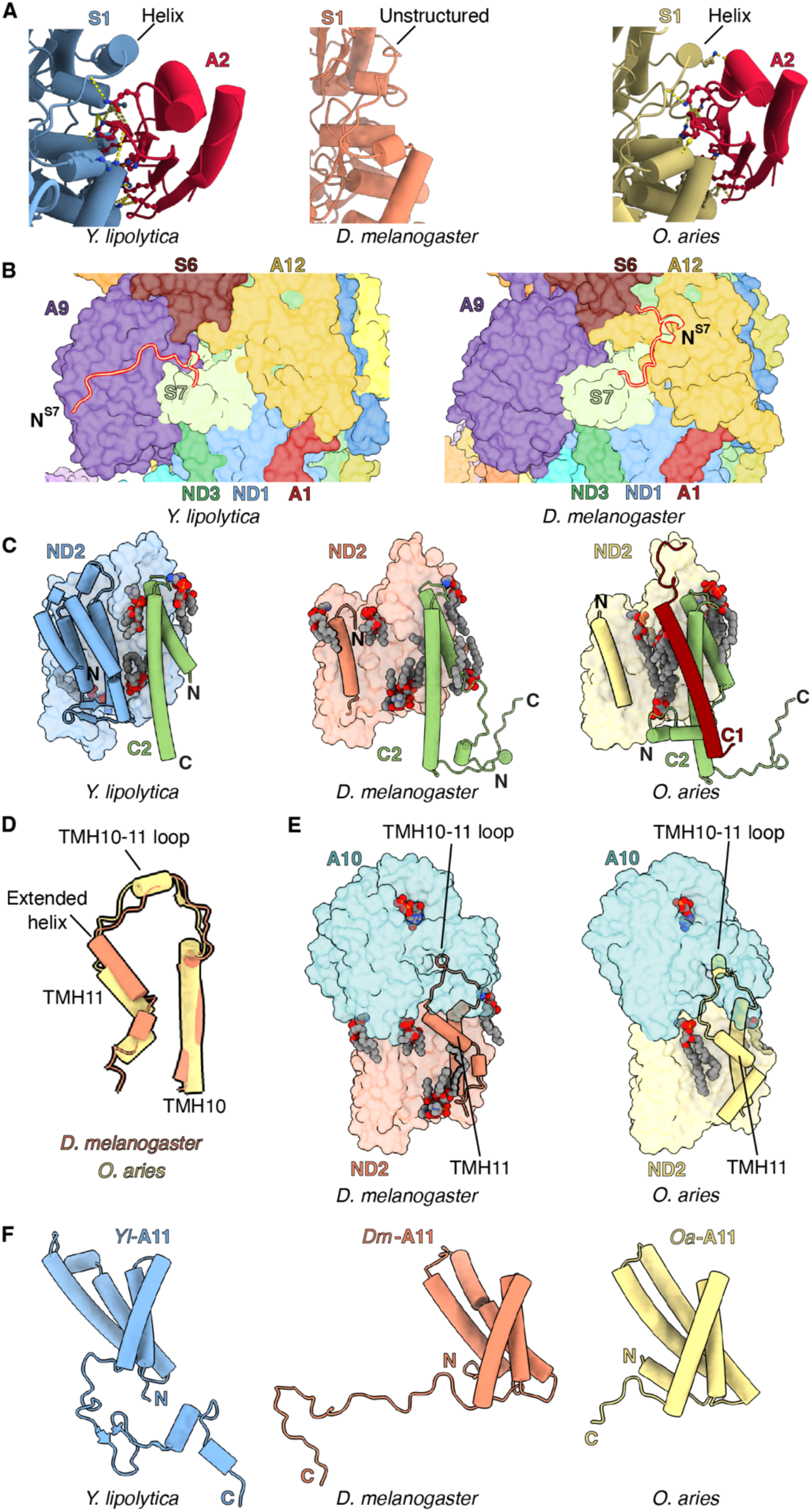
Features of *Dm*-CI subunits that would impact assembly and stability. (A) NDUFS1-NDUFA2 interface in *Y. lipolytic* (PDB: 6YJ4), *D. melanogaster* (this study) and *O. aries* (PDB :6ZKC) are shown. (B) The N-terminal extension of NDUFS7 in *Y. lipolytica* (PDB:6YJ4) and *D. melanogaster* are shown as cartoons. The other subunits are shown in surface colored as in Figure 1. (C) *Y. lipolytic* (PDB: 6YJ4), *D. melanogaster* (this study) and *O. aries* (PDB :6ZKC) ND2 is shown in surface. The N-terminal helices of ND2 are show as cartoons. NDUFC2 and NDUFC1 are shown as cartoons. Lipids are shown as spheres colored by element. (D) TMH10^ND2^, TMH11^ND2^ and TMH10-11^ND2^ loop of *D. melanogaster* and *O. aries* (PDB:6ZKC) are shown as cartoons (E) ND2-NDUFA10 interface in *D. melanogaster* (this study) and *O. aries* (PDB :6ZKC) is shown. ND2, NDUFA10 are shown in surface. TMH10^ND2^ TMH11^ND2^ and TMH10-11^ND2^ loop are shown as cartoons. Lipids are shown as spheres colored by element. (F) NDUFA11 in *Y. lipolytic* (PDB: 6YJ4) (blue), *D. melanogaster* (this study) (orange) and *O. aries* (PDB :6ZKC) (yellow) is shown as cartoons.

**Table 1.**
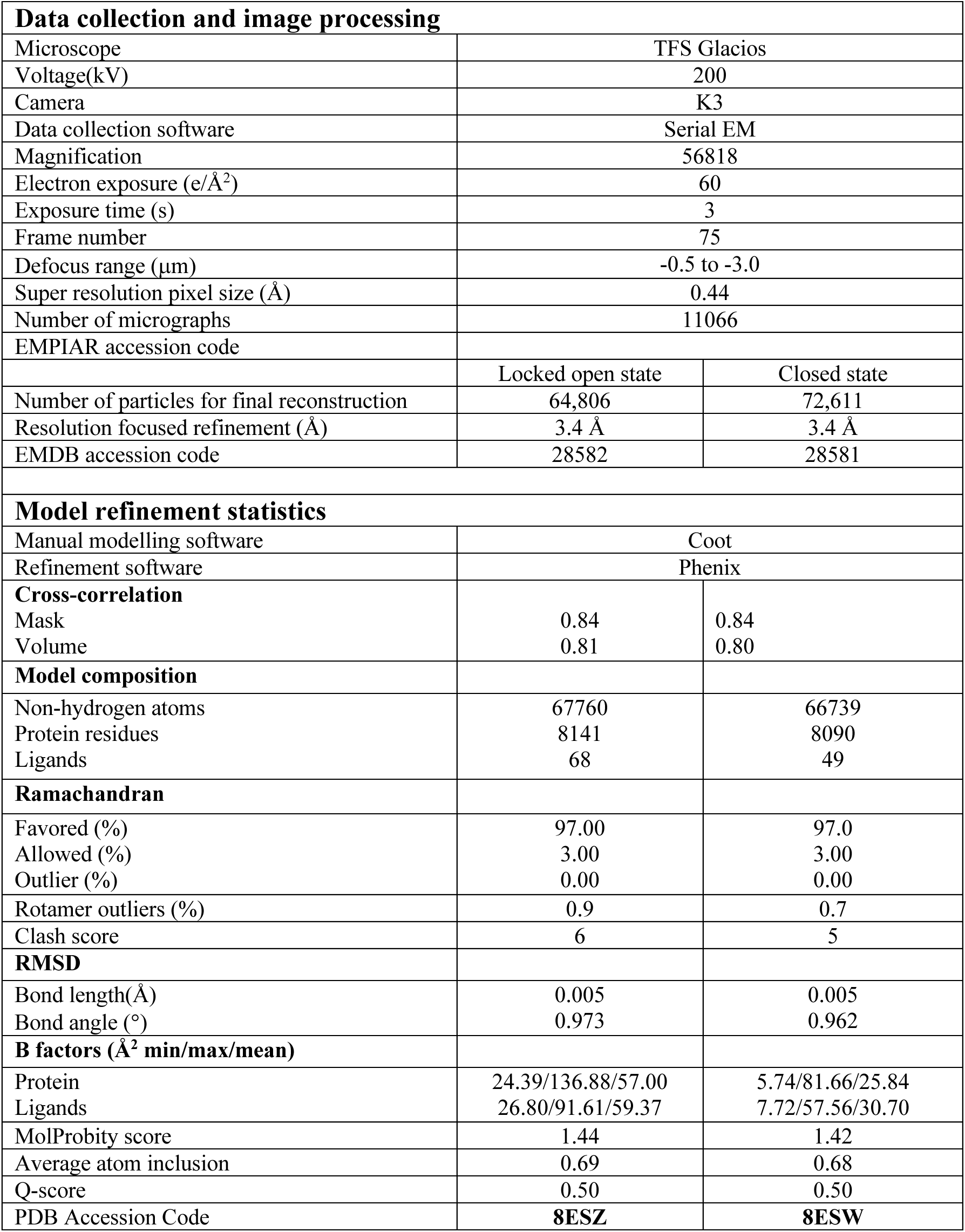

Our *Dm*-CI structure contained the 14 core subunits as well as 29 accessory subunits (Figure 1D and Figure 1-figure supplement 3), for a total composition of 43 subunits, two fewer than the 45 total subunits observed in mammals (Padavannil et al., 2022). Unlike *Y. lipolytica*, *T. thermophila* and plant CI, there were no accessory subunits unique to *Dm*-CI. An accessory subunit consistent with the position of NDUFV3 in mammals was present at sub-stoichiometric levels (see below). The electron transfer pathway from FMN to the final N2 cluster was conserved (Figure 1-figure supplement 4A and B). Although no quinone was added to the CI preparation, we were able to build a quinone molecule into density in the Q-tunnel (Figure 1-figure supplement 4C). The E-channel and the hydrophilic axis residues that are key to the coupling of electron transfer to the proton pumping were also conserved (Figure 1-figure supplement 4A and H) (Baradaran et al., 2013).

**Figure 3.**
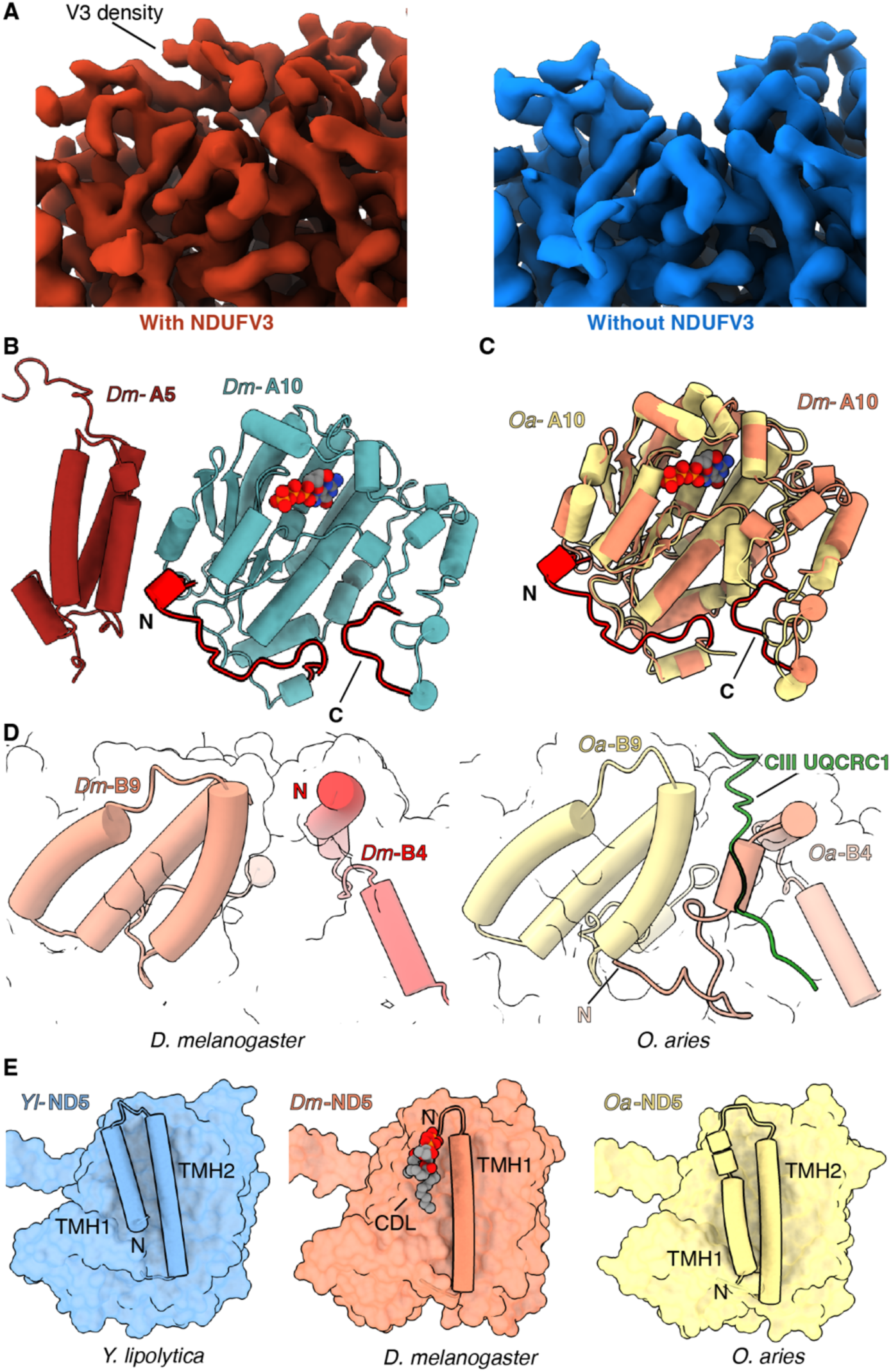
Features of *Dm*-CI subunits that may impact regulation, SC formation or lipid binding. (A) Cryo-EM map with (red) and without (blue) NDUFV3 density is shown. (B) NDUFA10-NDUFA5 interface in *Dm*-CI is shown. The subunits are shown in cartoons colored as in Figure 1. The N and C terminal extension of *Dm*-NDUFA10 are colored red.(C) Structural alignment of NDUFA10 from *O. aries* (yellow) (6ZKC) and *D. melanogaster* (orange) is shown. The N and C-terminal extensions of *Dm*-NDUFA10 are shown in red. (D) N-terminal region of NDUFB4 and NDUFB9 in *D. melanogaster* (this study) and *O. aries* (PDB:6QC3) is shown as cartoons. Loop region of *Oa*-CIII subunit UQCRC1 forming interface with *Oa*-CI subunit is show as cartoon colored in green (E) ND5 in *Y. lipolytic* (PDB: 6YJ4), *D. melanogaster* (this study) and *O. aries* (PDB :6ZKC) is shown as surface. The N-terminal helices of ND5 are show as cartoons. Lipids are shown as spheres colored by the element.

**Figure 4.**
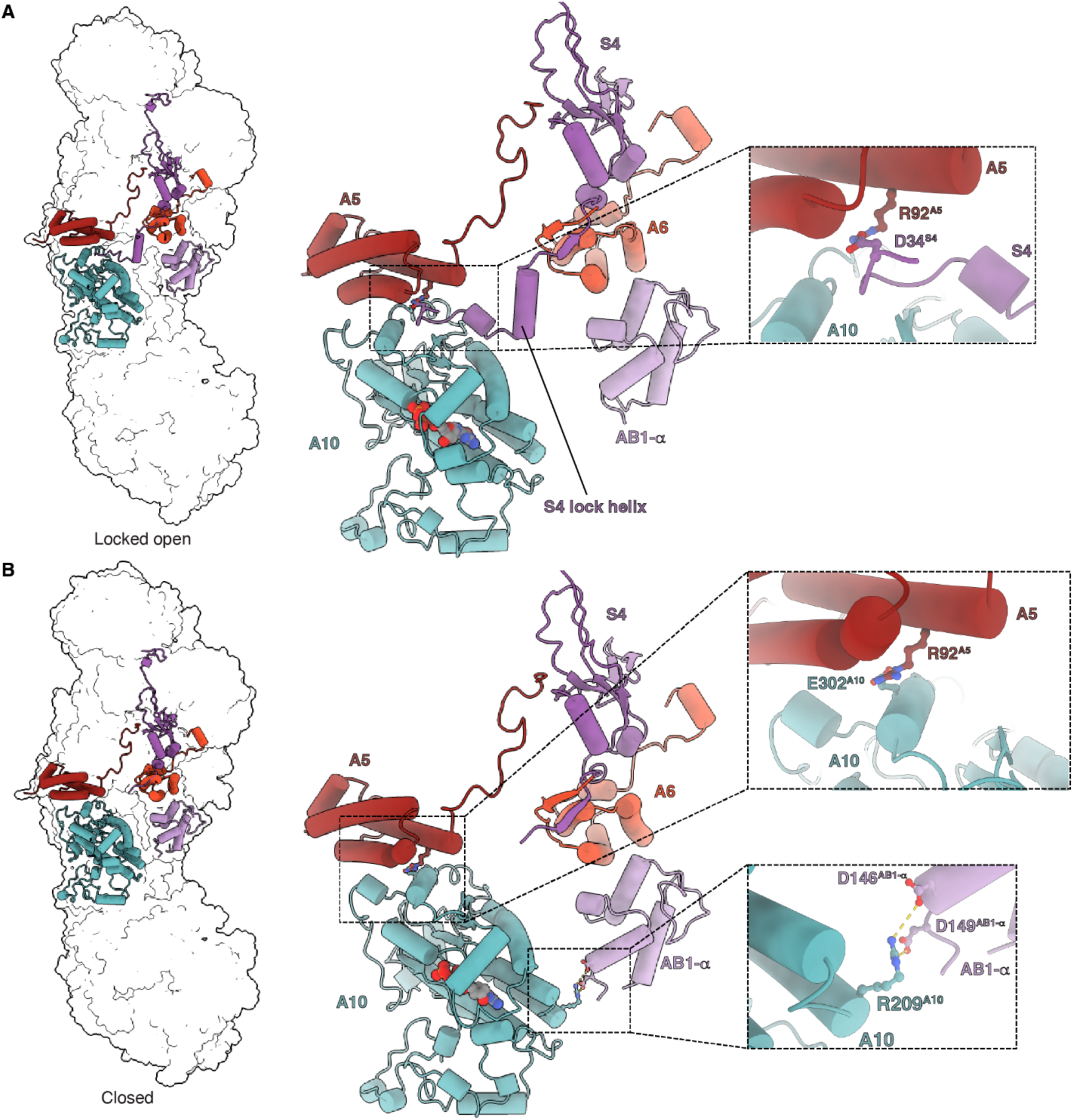
Matrix interactions in the locked open and closed states of *Dm*-CI. (A) *Dm*-CI subunits forming the bridge at the PA/MA interface in the locked open state are shown as cartoons colored as in Figure 1. The interaction between NDUFA5 and the N-terminal region of NDUFS4 is shown in the inset. (B) *Dm*-CI subunits forming the hinge at the PA/MA interface in the closed state are shown as cartoons colored as in Figure 1. NDUFA5, NDUFA10 and NDUFA10, NDUFAB1 interactions are show in insets.

The metazoan-specific transmembrane (TM) accessory subunit NDUFC1 is absent from the *Dm*-CI structure, consistent with the lack of a known ortholog (Garcia et al., 2017). Conversely, the N-module subunit NDUFA2, which is seen in all other known eukaryotic CI structures (Fiedorczuk et al., 2016; Maldonado et al., 2020; Parey et al., 2019; Zhou et al., 2022), is missing from the *Dm*-CI structure. A *D. melanogaster* ortholog of NDUFA2 (CG15434), which has a thioredoxin fold, co-migrates with CI on BN-PAGE gels (Garcia et al., 2017) (Figure 1-figure supplement 5A and B). The *Dm*-NDUFA2 ortholog was also seen in higher molecular weight bands that did not appear in immunoblots against *Dm*-CI core subunits, suggesting that *Dm*-NDUFA2 may be a component of multiple distinct complexes (Figure 1-figure supplement 5A and B). Given that a *Dm*-NDUFA2 band consistent with fully assembled *Dm*-CI was observed (Figure 1-figure supplement 5A and B), the lack of *Dm*-NDUFA2 in the structure indicates that it binds to the complex with lower affinity compared to other species and is lost during CI isolation (see further discussion below).

**Figure 5.**
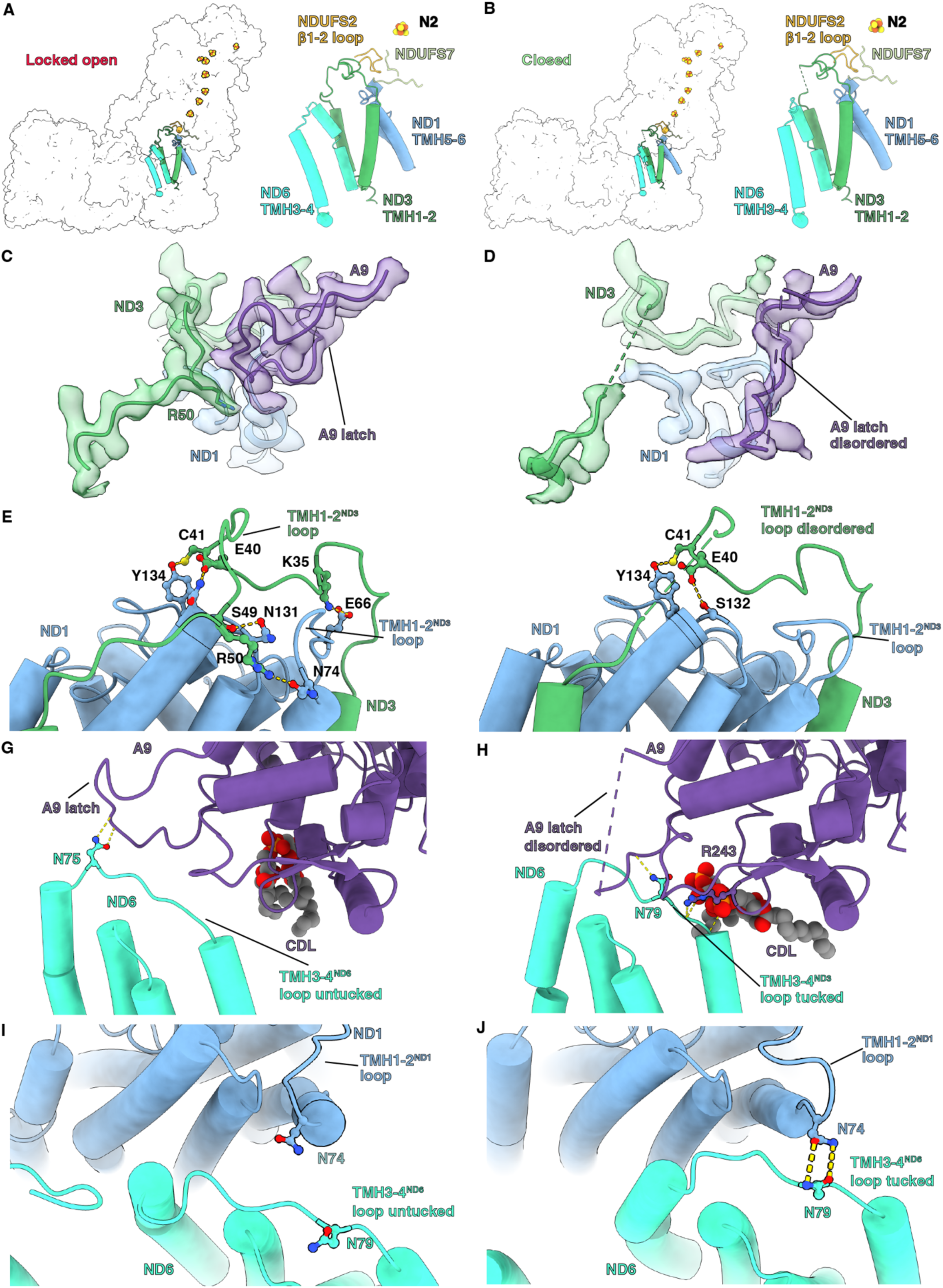
Q-site adjacent loops at the PA/MA interface. (A, B) The Q-site adjacent loops in the (A) locked-open state and (B) closed state are shown as cartoon colored as in Figure 1. (C, D) Interface between ND3, ND1 and NDUFA9 in the (C) locked-open and (D) closed state are shown. ND3, ND1 and NDUFA9 are shown as colored cartoons embedded in density colored as in Figure 1. (E, F) ND1 interaction with TMH1-2^ND3^ loop in the (E) locked-open and (F) closed states are shown. ND1 and ND3 are shown as cartoons colored as in Figure 1. (G, H) TMH3-4^ND6^ loop interaction with NDUFA9 in the (G) locked-open and (H) closed states are shown. ND6 and NDUFA9 are shown as cartoons colored as in Figure 1. (I, J) Interactions at the interface of ND1 and ND6 in the (I) locked-open and (J) closed states are shown. ND1 and ND6 are shown as cartoons colored as in Figure 1.

An ortholog of accessory subunit NDUFA3 was not annotated in the *D. melanogaster* proteome (Garcia et al., 2017). However, we identified density consistent with the presence of an NDUFA3 ortholog (Figure 1D, Video 2 and Figure 1-figure supplement 6). Amino acid assignment based on the observed side chain density allowed for the identification of an uncharacterized protein Dme1_CG9034, isoform B as the most likely NDUFA3 ortholog in *D. melanogaster* (Figure 1-figure supplement 6A and B). Consistent with this observation, proteomic analysis of a *Dm*-CI band cut from blue native gels identified CG9034 as one of the proteins that co-migrates with *Dm*-CI (Garcia et al., 2017); however, because it was not annotated and bioinformatics searches failed to identify it as an NDUFA3 ortholog, it was not identified as such.

While there were notable differences in the *Dm*-NDUFS1, *Dm*-NDUFS7, *Dm*-ND2 and *Dm*-ND5 structures (discussed below), the *Dm*-CI core subunits were overall like the core subunits of other opisthokonts (Figure 1-figure supplement 7). Further, differences in accessory subunits NDUFA11, NDUFC2, NDUFA10, NDUFB4 and NDUFB9 suggest differences in the assembly and regulation of *Dm*-CI, suggest how accessory subunits influence CI in mammals and reveal why *D. melanogaster* mitochondria have less supercomplex formation than mammals.

### Features of *Dm*-CI subunits with implications for assembly and stability

Differences between *Dm*-CI subunits and those of fungi and mammals suggest changes in the assembly and stability of the complex. For example, whereas the loop formed by amino acid residues 660-687 of NDUFS1 forms an α-helix in other species, in *Dm*-CI it is poorly resolved coil lacking secondary structure (Figure 2A, Video 3 and Figure 2-figure supplement 1A). This region forms part of the interface with accessory subunit NDUFA2 in other species, and the loss of the α-helix may be responsible for the absence of NDUFA2 in the *Dm*-CI structure (Figure 2-figure supplement 1C). Sequence alignment revealed that this loop is four residues shorter compared to *Y. lipolytica* and mammals and that several bulky residues have been replaced by alanine or glycine residues in *Dm*-NDUFS1 (Figure 2-figure supplement 1D) resulting in the loss of ordered secondary structure and increased flexibility. The helical structure in the NDUFS1 loop is not seen in bacterial CIs (Baradaran et al., 2013; Kolata and Efremov, 2021; Kravchuk et al., 2022), suggesting that this secondary structure element may have evolved in eukaryotes specifically to interact with NDUFA2.

In mammals, NDUFA2 plays an important role in the stability of the N-module (Stroud et al., 2016) and has been proposed to play a role in the regulation of CI by ROS (Padavannil et al., 2022). Whereas NDUFA2 knockout in HEK293T cells results in the loss of the N-module and complete loss of CI activity (Stroud et al., 2016), NDUFA2 knock-down in *D. melanogaster* has minimal effects, with CI retaining ∼97% of WT (Mhc-Gal4/W^1118^ flies) activity (Garcia et al., 2017), indicating that NDUFA2 is not needed for *Dm*-CI assembly or stability. Given that NDUFA2 only interacts with NDUFS1, it was proposed that NDUFA2 binding compensates for truncation of NDUFS1’s “D domain” that is otherwise present in bacterial orthologs (Figure 2-figure supplement 1B and C) (Padavannil et al., 2022). However, like other eukaryotes, domain D remains short in *Dm*-NDUFS1 and the reason for the sustained stability of *Dm*-CI in the absence of NDUFA2 is unclear. Further work is needed to understand why *Dm*-CI assembly and stability appears independent of *Dm*-NDUFA2.

As in *Y. lipolytica*, NDUFS7 in *D. melanogaster* has an extended N-terminus relative to mammalian CI (Figure 2B, Figure 2-figure supplement 2A and B). However, whereas the N-terminus of *Yl*-NDUFS7 binds along the surface of NDUFA9, that of *Dm*-CI is flipped ∼180° binding overtop of NDUFA12 (Figure 2B, Figure 2-figure supplement 2A). This additional interface would stabilize the association of NDUFA12 in *Dm*-CI, which has implications for assembly of this part of the complex. During CI assembly, the assembly factor NDUFAF2 binds at the equivalent position of NDUFA12 (Parey et al., 2019) and is exchanged for NDUFA12 before full assembly of the PA (Andrews et al., 2013; Vogel et al., 2007). The additional interactions between NDUFS7 and NDUFA12 in *Dm*-CI may thus influence assembly of the PA by promoting the exchange of NDUFAF2 with NDUFA12 through stabilization of NDUFA12 binding (Figure 2B).

As in the mammalian CI, *Dm*-ND2 lacks three TMHs at the N-terminus, thus having 11 TMHs as opposed to the 14 TMHs otherwise seen in bacteria, plants, ciliates, and yeast (Birrell and Hirst, 2010) (Figure 2C and Figure 2-figure supplement 3A). The cavity formed by the lack of the three TMHs is filled with lipids that are held in place in part by the NDUFC2 subunit (Figure 2C). Similar to other eukaryotic CIs, the last TMH in *Dm*-ND2 (TMH11^ND2^), has two additional turns, compared to TMH11^ND2^ of mammals (Figure 2D and Figure 2-figure supplement 3B) (Klusch et al., 2021; Parey et al., 2019; Zhou et al., 2022). It has been proposed that in mammals subunit NDUFC1 plays a role in shortening TMH11^ND2^ by binding a cardiolipin molecule that caps the helix stabilizing its partially unwound state (Padavannil et al., 2022). Given that *D. melanogaster* lacks accessory subunit NDUFC1 and has a longer TMH11^ND2^ relative to mammals, this supports the proposed role of NDUFC1 in mammals. Lack of NDUFC1 in *Dm*-CI also indicates that it was recruited as an accessory subunit only after the split of Protostomia and Deuterostomia.

In both mammals and *D. melanogaster*, the TMH10-11^ND2^ loop provides a major interface with metazoan-specific subunit NDUFA10 (Figure 2E and Video 4). In *Y. lipolytica*, which lacks any subunit binding on the matrix side of ND2, as well as in plants and Tetrahymena which use the equivalent loop to interact with their γ-carbonic anhydrase subunit (Klusch et al., 2021; Maldonado et al., 2020; Soufari et al., 2020; Zhou et al., 2022), the TMH10-11^ND2^ loop spans across the matrix surface of ND2 as a coil. In mammals, TMH10-11^ND2^ spans the same distance across the matrix surface but forms a short α-helix that interacts directly with NDUFA10 (Figure 2D and E and Video 4). Given that the length of the TMH10-11^ND2^ loop is only shorter by two residues in *D. melanogaster* compared to mammals (Fig.2-figure supplement 3B), the additional residues required for the TMH10-11^ND2^ loop to fold into an α-helix in mammals must come from the unwinding of TMH11^ND2^. Thus, a simple model emerges for how mammalian CI is dependent on NDUFC1 for assembly. Namely, the mammalian interface between ND2 and NDUFA10 cannot form before NDUFC1 binds and recruits a cardiolipin to partially unwind TMH11^ND2^ (Figure 2-figure supplement 3C) (Padavannil et al., 2022). This model is consistent with the known binding order and dependencies of NDUFC1 and NDUFA10 to ND2 during CI assembly (Guerrero-Castillo et al., 2017; Stroud et al., 2016).

In *Dm*-CI, due to an extended C-terminal coil, the MA accessory subunit NDUFA11 has a much more extensive interaction with the core subunits than seen in mammals (Figure 2F, Figure 2-figure supplement 4A and 4B and Video 5). NDUFA11 is a four-TMH subunit that binds adjacent to ND2 atop the ND5 lateral helix (ND5-HL) and ND5 TMH15 (TMH16 in fungi and mammals). It has an arch shape with the concave surface facing CI. The cavity formed at the interface is filled with lipids which bridge between NDUFA11 and ND2/ND5. In mammalian CI, only limited protein-protein contacts between NDUFA11 and other CI subunits are observed, dominated mainly by its short C-terminal coil (Figure 2F and Figure 2-figure supplement 4A). For this reason, detergent extraction can result in the loss of NDUFA11 in mammals, resulting in so-called ‘state 3’ particles (Chung et al., 2022b; Fiedorczuk et al., 2016; Zhu et al., 2016). However, in the fungus *Y. lipolytica* the C-terminus of NDUFA11 is much longer and extends along the matrix side of the complex (Parey et al., 2019). Likely for this reason NDUFA11 in *Y. lipolytica* is harder to dissociate from the complex even after treatment with harsh detergents (Angerer et al., 2011).

Like *Y. lipolytica*, NDUFA11 in *Dm*-CI has an extended C-terminus (Figure 2F and Figure 2-figure supplement 4A). However, unlike the C-terminus in *Y. lipolytica* NDUFA11 which runs along the length of the MA, that of *Dm*-NDUFA11 runs across the membrane arm (i.e., perpendicular to the long axis), tucking between NDUFB5, NDUFS5, NDUFA8 and ND2 and emerging adjacent to NDUFC2 on the opposite side (Figure 2F and Video 5). This arrangement is not seen in any other know CI structure. The C-terminus of *Dm*-NDUFA11 occupies space that is occupied by subunit NDUX1 (NUXM) in yeast, plants and ciliates (a subunit that is lost in metazoans along with the truncation of ND2) and the N-terminus of NDUFC2 in mammals (Fiedorczuk et al., 2016; Klusch et al., 2021; Maldonado et al., 2020; Parey et al., 2019; Zhou et al., 2022). The extended C-terminus of *Dm*-NDUFA11 has significant implications for the assembly of *Dm*-CI. Mammalian CI is assembled through a series of intermediates and NDUFA11 is a terminally associated protein that does not form a part of any assembly intermediate (Guerrero-Castillo et al., 2017). Also in mammals, NDUFB5, NDUFB8 and NDUFS5 are all present in assembly intermediates prior to the addition of NDUFA11 (Guerrero-Castillo et al., 2017). The arrangement seen in *Dm*-CI, in which the C-terminus of *Dm*-NDUFA11 is sandwiched between *Dm*-NDUFB5, *Dm*-NDUFA8, *Dm*-NDUFS5 and *Dm*-ND2 (Video 5), suggests that *Dm*-NDUFA11 would need to bind *Dm*-ND2 before *Dm*-NDUFB5, *Dm*-NDUFA8 and *Dm*-NDUFS5. However, as NDUFA11 binds on top of ND5-HL, ND5 would need to associate with ND2 before NDUA11 could bind. This order of events is counter to what occurs in mammals (Guerrero-Castillo et al., 2017) and indicates that, like plant CI (Ligas et al., 2018), assembly of this region of *Dm*-CI might proceed via a distinct mechanism than that of mammals.

### Features of *Dm*-CI subunits with implications for regulation

In mammals there are two isoforms of the NDFUV3 subunit, a long isoform NDUFV3-L and a short isoform NDUFV3-S and it has been proposed that binding of the different isoforms may impact the activity of CI (Bridges et al., 2016; Dibley et al., 2017; Guerrero-Castillo et al., 2016). Density for a subunit at the position of NDUFV3 in mammals was also observed in the *Dm*-CI structure (Figure 3A). However, in the average structure calculated using all *Dm*-CI particles this density was weak compared to that of the surrounding core subunits. When focused refinements were performed with a mask around the tip of the PA, two clear classes could be isolated differing in the occupancy of the NDUFV3 site (Figure 3A and Figure 3-figure supplement 1). Thus, in *Dm*-CI this site is only partially occupied.

The 68 kDa fragment of the atypical cadherin (Ft4) can regulate *Dm*-CI activity (Sing et al., 2014) and it was proposed that it may bind to *Dm*-CI at the NDUFV3 site (Bridges et al., 2016). The density of the occupied class was too noisy to confidently assign the sequence of the subunit from the map (Figure 3A). However, our results are consistent with the hypothesis that binding at this site may be regulatory (Bridges et al., 2016). One possible explanation for the noisy density and inability to fully classify the particles completely (Figure 3-figure supplement 1) is that different proteins may be bound at this site potentially allowing for more complex regulation.

In addition to the major interface between NDUFA10 and ND2, in mammals NDUFA10 forms a state dependent interface with accessory subunit NDUFA5 (Figure 3-figure supplement 2B) (Agip et al., 2018; Kampjut and Sazanov, 2020; Letts et al., 2019). In *Dm*-CI, the interface between NDUFA10 and NDUFA5 is larger due to an extended NDUFA10 N-terminal coil that inserts between NDUFA10 and NDUFA5 (Figure 3B and C and Figure 3-figure supplement 2A). It has been debated whether the breaking of the interface between NDUFA10 and NDUFA5 occurs during enzyme turnover or is a feature of the D state (Agip et al., 2018; Kampjut and Sazanov, 2020). The enhanced interface between *Dm*-NDUFA10 and *Dm*-NDUFA5 would make these subunits more difficult to separate and indicates that, although the interactions at this interface may be state dependent (see below), the interface is less likely to be fully broken as seen in the mammalian context (Agip et al., 2018; Kampjut and Sazanov, 2020; Letts et al., 2019).

### Features of *Dm*-CI with implications for SC assembly

The structure provides a basis for understanding the lower abundance of SCs between *D. melanogaster* and mammals. In mammals, the N-terminus of NDUFB4 forms part of the only matrix interface between CI and CIII_2_ in SC I+III_2_ (Letts et al., 2019; Letts and Sazanov, 2017). In this interaction a loop from the UQRC1 subunit of one CIII protomer binds in between the N-terminus of NDUFB4 and the three-helix-bundle of subunit NDUFB9. In *Dm*-CI the N-terminus of *Dm*-NDUFB4 is truncated relative to that of mammals and does not extend far enough towards NDUFB9 to form this interface (Figure 3D and Figure 3-figure supplement 3). Thus, the lack of this interface likely contributes to the observed low abundance of SCs in *D. melanogaster* mitochondria.

Additionally, given its role in bridging between CI and CIII_2_ in respiratory SCs (Letts et al., 2019; Letts and Sazanov, 2017), it has been proposed that NDUFA11’s interaction with CI may have been weakened in species, such as mammals, to promote SC formation by requiring the presence of CIII_2_ to hold NDUFA11 in place (Padavannil et al., 2022). The extended interaction interface provided by the C-terminus of *Dm*-NDUFA11 indicates that *Dm*-NDUFA11 is not dependent on the presence of CIII_2_ for its stable association with the *Dm*-CI. Thus, the *Dm*-CI structure supports the proposed inverse relationship between the strength of the interaction between NDUFA11 and CI and the abundance of SCs (Padavannil et al., 2022).

### Differences in lipid binding

Lipids from an integral part of the CI MA and loss of lipids during purification results in diminished CI activity (Letts et al., 2016; Padavannil et al., 2022). Across species structural lipids are seen tightly binding to the surface of CI at the interface of the core H^+^-pumping subunits and specific deformation of the lipid membrane by NDUFA9 occurs adjacent to the Q-tunnel (Kampjut and Sazanov, 2020; Parey et al., 2019; Zhou et al., 2022). In general, the pattern of lipid binding to *Dm*-CI (Figure 1-figure supplement 4I) is like that seen in other CI structures with two notable exceptions. First, instead of 16 TMHs observed in ND5 of *Y. lipolytica* and mammals, *Dm*-ND5 lacks the first TMH for a total of only 15 (Figure 3E and Figure 3-figure supplement 4A). Like what is seen with the metazoan specific shortening of ND2 discussed above, the region occupied by ND5-TMH1 in other species is not occupied by any other protein subunit but binds lipid (Figure 3E, Figure 3-figure supplement 4A). As mammalian ND5 maintains its full complement of helices, loss of the first TMH must have occurred after the split of Protostomia and Deuterostomia, but how widespread the ND5 deletion is in Protostomia along with any functional implications remains to be determined.

Second, in mammals the N-terminus of the two-TMH accessory subunit NDUFC2 binds underneath ND2 in the pocket left by the deletion of the first three ND2 TMHs. However, in *Dm*-CI this space is filled by the C-terminus of NDUFA11 (Figure 2C and Video 5). Like mammalian NDUFC2, *Dm*-NDUFC2 has an extended N-terminus relative to that of *Y. lipolytica* (Figure 2C and Figure 3-figure supplement 5A), however, instead of crossing over TMH2^C2^ and binding under ND2, it crosses over the C-terminal coil of NDUFA8 and forms additional interactions with NDUFB1 and NDUFB5 (Figure 2C and Figure 3-figure supplement 5A). These additional interactions contribute to lipid binding at the ND2/ND4 interface by capping this pocket and likely help to stabilize lipid binding at the interface between the two H^+^-pumping core subunits.

### Focused classification of Dm-CI reveals a NDUFS4 helix-locked state

Initial poor resolvability of the average *Dm*-CI map around NDUFA10 at the interface of the PA and MA led us to perform focused classifications using a mask encompassing subunits NDUFA10, NDUFA5, NDUFA6 and NDUFAB1-α (Figure 4-figure supplement 1). This classification revealed two major classes and two minor classes of *Dm*-CI particles that differ in the presence of an ordered α-helical element from NDUFS4 bound at the interface of NDUFA5, NDUFA10 and NDUFA6 and in the angle between the PA and MA (Figure 4). We call the two major states the NDUFS4-helix-locked open (locked open) state and the closed state (Figure 4). Consistent with what has been described in mammals, *Y. lipolytica* and bacteria (Kampjut and Sazanov, 2020; Kravchuk et al., 2022; Parey et al., 2021), in this naming scheme “open” and “closed” refer to the angle between the PA and the MA, with the open state angle being larger (Figure 4-figure supplement 2A). Like the major closed state, the two minor classes lacked the NDUFS4 helix and differed from the major closed state only by the angle between the PA and MA (Figure 4-figure supplement 1). Unlike what is seen in other CI structures the open state of *Dm*-CI is accompanied by the insertion of an N-terminal helix from NDUFS4 as a wedge between the PA and the MA (Figure 4A). Three-dimensional variability analysis (3DVA) performed on the full set of particles revealed a smooth transition between the two states along with the disappearance of the NDUFS4 helical density (Video 6). This indicates that the vitrified *Dm*-CI particles can adopt conformations along the trajectory from the major closed state to the open state and once in the open state they can be “locked” open by the binding of the NDUFS4 helix (Figure 4A and Video 6). The N-terminal region of NDUFS4 that forms the lock helix is not conserved in *Y. lipolytica* or mammals (Figure 4-figure supplement 3) suggesting that it evolved after the split of Protostomia and Deuterostomia.

The angle between the MA and the PA in the helix-locked open state is most similar to the angle in the open states of *Y. lipolytica* and mammals (Figure 4-figure supplement 4). This is also the case when comparing the *Dm*-CI closed state to that of other organisms (Figure 4-figure supplement 4). Although the density of the NDUFAB1-α subunit is weak due to flexibility, its position in the close state is consistent with the formation of salt-bridges between residues D146^AB1-α^, D149^AB1-α^ and R209^A10^ (Figure 4B and Video 6). In plant and Tetrahymena CI, the NDUFAB1-α subunit is involved in bridging interactions between the PA and MA (Klusch et al., 2021; Zhou et al., 2022); however, to our knowledge, this is the first indication of direct bridging between the PA and MA via NDUFA10 and NDUFAB1-α in any opisthokont structure.

Although NDUFA5 and NDUFA10 remain in direct contact in both the locked open and the closed states, there is a state-dependent change in their interaction (Figure 4 and Video 6). In the open state, the interaction between NDUFA5 and NDUFA10 is mediated in part by the N-terminus of NDUFS4 which binds along the surface of NDUFA10 and D34^S4^ forms a salt bridge with R92^A5^ (Figure 4A and Video 6). In the closed state, the N-terminus of NDUFS4 is disordered and NDUFA5 slides along the surface of NDUFA10 such that R92^A5^ forms a salt bridge with E302^A10^ (Figure 4B and Video 6). Thus, R92^A5^ is used to form salt bridging interactions with NDUFS4 and NDUFA10 in a state-dependent manner (Figure 4 and Video 6).

### In the helix-locked open state the CoQ reduction site loops are buried by an NDUFA9 “latch”

Conformational changes in loops adjacent to the CoQ reduction site were seen between the locked open and closed states of *Dm*-CI (Figure 5 and Video 7). In mammals, *Y. lipolytica* and *E. coli* the open state of the complex is commonly associated with specific conformations or disorder of loops around the CoQ binding site (α1-2^S7^ loop, α2-ý1^S7^ loop, TMH5-6^ND1^ loop, TMH1-2^ND3^ loop, β1-2^S2^ loop and TMH3-4^ND6^ loop) as well as a π-budge in TMH3^ND6^ (Kampjut and Sazanov, 2020; Kravchuk et al., 2022; Parey et al., 2021, 2018). In the closed state these loops are generally well ordered and TMH3^ND6^ re-folds into an α-helix (Figure 5-figure supplement 1). The open and closed states of *Dm-*CI do not follow these trends (Figure 5-figure supplement 1). No differences were seen in *Dm*-CI between the locked open and closed states in the conformations of the α1-2^S7^, TMH5-6^ND1^ and β1-2^S2^ loops (Figure 5-figure supplement 1A, 1B and 1C). The TMH1-2^ND3^ loop and TMH3^ND6^ showed the opposite trend compared to other species: the TMH1-2^ND3^ loop is well ordered in the locked open state and partially disordered in the closed state; and TMH3^ND6^ is α-helical in the open state and contains a π-budge in the closed state (Figure 5A and B and Figure 5-figure supplement 1D and 1E). These differences in the TMH1-2^ND3^ loop stem from a state-dependent interaction with the C-terminal loop of NDUFA9, a rotation of ND1 relative to the rest of the MA and the movement of TMH4^ND6^ and the TMH3-4^ND6^ loop (Figure 5C-J Video 7, and Figure 5-figure supplements 1 and 2).

In the locked open state, the C-terminal loop of NDUFA9 binds atop the TMH1-2^ND3^ loop, trapping R50^ND3^ to the surface of ND1 (Figure 5C), thereby holding the loop in place like a latch. In the closed state the C-terminal loop of NDUFA9 moves away from the TMH1-2^ND3^ loop, providing space for conformational flexibility and both loops become partially disordered (Figure 5D). Specifically, clear density is lost for R50^ND3^ and surrounding residues (Figure 5D). The movement of NDUFA9 away from the TMH1-2^ND3^ loop is accompanied by changes in the interactions between ND1 and the TMH1-2^ND3^ loop caused by the rotation of ND1 and conformational changes in the TMH1-2^ND1^ loop. In the locked open conformation multiple hydrogen bonding and salt bridging interactions were seen between the TMH1-2^ND3^ loop and ND1 (Figure 5E). However, in the closed state most of these interactions are lost except for a hydrogen bond between the conserved C41^ND3^ and Y134^ND1^ (Figure 5E and F). E40^ND3^ swaps hydrogen bonding partners from N133^ND1^ in the locked open state to S132^ND1^ in the closed state (Figure 5E and F). Fewer hydrogen bonds between ND1 and the TMH1-2^ND3^ loop would also contribute to the observed higher flexibility of this loop in the closed state (Figure 5C, D, E and F). Finally, the TMH3-4^ND6^ loop moves relative to NDUFA9, going from exposed or “untucked” in the locked open state to “tucked” under NDUFA9 in the closed state (Figure 5G and H and Video 7). The movement of TMH3-4^ND6^ loop brings it into contact with the TMH1-2^ND1^ loop in the closed state and requires an ∼10 Å translation of TMH4^ND6^ (Figure 5I and J, Figure 5-figure supplement 2 and Video 7). Similar “latching” behavior of the NDUFA9 C-terminus and “tucking” of the TMH3-4^ND6^ loop, though different in the structural details, was also recently seen in the structure of the thermophilic yeast *Chaetomium thermophilum* (Laube et al., 2022), suggesting that this may be a general mechanism for regulating the activity.

## Discussion

We present here the first structure mitochondrial CI from the model organism *D. melanogaster*. This represents the first CI structure from an insect and from any protostomian, a broad group of animals that split from mammals and other deuterostomians ∼600 million years ago. During the evolution of metazoans, a split in the biochemical behavior of mitochondrial CI occurred corresponding to the Protostomia/Deuterostomia divide (Figure 1A). According to established biochemical assays (Babot et al., 2014), all characterized deuterostomian CIs can enter a deactive “D” state, a property which it shares with fungal CIs, whereas this is not the case for protostomian CIs (Figure 1A) (Babot et al., 2014; Maklashina et al., 2003). However, it is important to note that although all current data are consistent with a Protostomia/Deuterostomia split in the biochemical behavior of CI, only a small subset of species from each group have been investigated (Figure 1A). Characterization of additional species across both groups will provide a better understanding about the evolution of this functional split. Recently the structural basis of the A-to-D transition has become central to the debate over the CI coupling mechanism (Chung et al., 2022a; Kampjut and Sazanov, 2022) and the unique features of our *Dm*-CI structures inform this debate.

In fungi and mammals, the state-dependent accessibility of the ND3 cysteine in the active “A” and D states is clearly understood from CI structures. The yeast *Y. lipolytica* and mammalian structures have been solved in multiple states that broadly fall into two categories, either “open” or “closed”, defined by the angle between the PA and the MA (Agip et al., 2018; Kampjut and Sazanov, 2020; Parey et al., 2021). In the open states the TMH1-2^ND3^ loop, which harbors the reactive cysteine, is disordered indicating flexibility and accessibility, whereas in the closed state the TMH1-2^ND3^ loop is well ordered and the reactive cysteine is buried and inaccessible. This led to the proposal that the open state of CI corresponds to the D state and the closed state of the complex corresponds to the A state (Agip et al., 2018; Zhu et al., 2016). This proposal is supported by the structure of deactivated bovine CI which was found to contain 87.5% open state particles (Blaza et al., 2018). However, it was also found that under turnover conditions ovine CI adopts open states, leading Kampjut and Sazanov to propose that open states are part of the catalytic cycle and that the deactive state is a particular ‘deep’ open state (Kampjut and Sazanov, 2020). They test their model by deactivating the ovine complex prior to structure solution and reporting a large conformational shift in TMH4^ND6^ (Figure 5-figure supplement 2) (Kampjut and Sazanov, 2020). Thus, they conclude that the deactive state is a specific open state characterized by a large displacement of TMH4^ND6^ (Kampjut and Sazanov, 2022, 2020; Kravchuk et al., 2022). However, all deposited maps of the deactivated ovine complex (EMDB-11260, EMDB-11261, EMDB-11262 and EMDB-11263) show very weak or no density for ND5-HL, TMH16^ND5^, NDUFA11 or TMH4^ND6^ (Kampjut and Sazanov, 2020). Poor density in these regions have been associated with so-called “state 3” particles which are proposed to correspond to CI in the first stages of dissociation (Chung et al., 2022b; Zhu et al., 2016). This leads to the possibility that, despite being able to measure activity, the deactivation treatment of Kampjut and Sazanov may have partially denatured the ovine complex or sensitized it to disruption during the cryoEM grid preparation (Kampjut and Sazanov, 2020). If these structures represent the D state, then it follows that disorder of ND5-HL, TMH16^ND5^ and NDUFA11 in addition to the movement of TMH4^ND6^ would also be features of the D state. Although that is possible, despite lower resolution, these features were not observed in the deactivated bovine complex (EMDB-3731) (Blaza et al., 2018). Altogether, it is clear that CI’s deactive state is an open state. However, given the discrepancies between mammalian structures, it remains unclear how the deactive open state differs from other open states that may be part of CI’s catalytic cycle.

The proposed function of open states during catalysis are two-fold. Firstly, the disordering of the CoQ-site loops, in particular the TMH1-2^ND3^ and β1-2^S2^ loops, would disrupt the CoQ binding site and open the CoQ-binding cavity to solvent after the formation of ubiquinol (CoQH_2_), thereby “washing out” the highly hydrophobic substrate that may otherwise remain “stuck” in the active site tunnel. Secondly, an open state during turnover would disrupt the hydrophilic axis via the formation of a π-bulge in TMH3^ND6^, rotating hydrophobic residues into the axis and preventing a futile cycle caused from “back flow” of protons to the solvent accessible CoQ site (Kampjut and Sazanov, 2022, 2020; Kravchuk et al., 2022).

Alternatively, it has been proposed that that catalysis only occurs through a series of closed states and that all observed open states correspond to the D state (Chung et al., 2022a, 2022b). In this model, the multiple observed open states would stem from increased flexibility between the PA and MA when the angle between them increases and the TMH1-2^ND3^ loop is released from its binding site at the PA/MA interface. Therefore, CoQ entry and exit would occur though standard diffusion in and out of the active site tunnel with more subtle conformational changes resulting in changing affinities for CoQ and CoQH_2_ (Chung et al., 2022b, 2022a). Both models are consistent with conformational changes in the CoQ site loops being important during turnover, which has been demonstrated through site-specific cross-linking of the TMH1-2^ND3^ loop with NDUFS7 (Cabrera-Orefice et al., 2018), but differ in the degree of conformational change needed. Also, this closed-states-only model of turnover proposes that the α-helix-to-π-bulge transition of TMH3^ND6^ does not occur during catalysis and that the TMH3^ND6^ π-bulge is only a feature of the D state.

The structures of resting *Dm*-CI reported here are not fully consistent with either of these mechanistic models. First, in the helix-locked open state, the angle between the PA and MA are more consistent with the open states of *Y. lipolytica* and mammalian CIs but the active site loops are fully ordered and buried by the NDUFA9 latch (Figure 5C). This state also lacks the TMH3^ND6^ π-bulge indicating that the water wire of the hydrophilic access is intact, though a higher resolution structure is needed to confirm this. Second, only in the closed state, as defined by the angle in between the PA and MA, did we see increased flexibility in the CoQ site loops and the formation of the TMH3^ND6^ π-bulge.

Given that both helix-locked open and closed states are resting-state structures, i.e., no substrate was provided to the complex prior to vitrification, we cannot make strong conclusions regarding the catalytic relevance of these states. However, conformational changes in the TMH1-2^ND3^ loop are necessary for the coupling between CoQ reduction and H^+^-pumping (Cabrera-Orefice et al., 2018). Therefore, it is more likely that the closed state in which this loop is freer to undergo conformational changes is an on-pathway state and that the helix-locked open state is an off-pathway resting state, similar in function but biochemically distinct from the D state of yeast and mammals. Flexibility and opening of the TMH1-2^ND3^ loop during turnover are also supported by the enhanced reactivity of the ND3 cysteine seen in our 2 mM NEM pre-activation conditions (Figure 1B). Given that the proposed physiological function of the D state is to prevent reactive oxygen species production by reverse electron transport (Chouchani et al., 2013), it is possible that multiple distinct mechanisms have evolved to achieve this end. If the helix-locked state is a D-like state, it suggests that reverse electron transport by CI can be blocked in two distinct ways: 1) opening the CoQ site to solvent and thereby preventing CoQH_2_ binding as seen in the standard D state; or 2) blocking conformational changes needed for coupling between CoQ reduction/CoQH_2_ oxidation and H^+^-pumping as seen in the NDUFA9-latched-helix-locked-state. The presence of NDUFA9-latch holding the TMH1-2^ND3^ loop in place also explains the insensitivity of this potential D-like state to NEM (Figure 1B).

If the *Dm*-CI closed state is a catalytically competent state it supports the hypothesis that the TMH3^ND6^ π-bulge forms as part of the catalytic cycle as it is the closed state of *Dm*-CI that displays the TMH3^ND6^ π-bulge structure. This interpretation also suggests that what matters is not the angle between the PA and MA but whether the active site loops are free to undergo conformational changes. Thus, it is likely that CI can turnover without large changes in the angle between the PA and MA. This would be consistent with species whose CI have additional bridging interactions between the PA and MA that may limit changes in the PA/MA angle, such as plants, *Tetrahymena* and the thermophilic yeast *Chaetomium thermophilum* (Klusch et al., 2021; Laube et al., 2022; Zhou et al., 2022).

Another possible interpretation of the two states would be that they are both resting states associated with distinct phases of catalysis, i.e., they are both on-pathway states. However, this is unlikely as the NDUFS4 lock-helix is not universally conserved nor adjacent to the active site (Figure 4-figure supplement 3). Therefore, it is more likely that the NDUFS4 lock-helix is a regulatory element that has evolved in a specific branch of the eukaryotic tree. If the helix-locked state is an auto-inhibited state, these structures represent a novel regulatory mechanism that may be exploited to inhibit CI turnover in other species.

Recently, structures of *E. coli* CI, which does not deactivate (Maklashina et al., 2003), revealed a variety of open states (Kravchuk et al., 2022). In this study, a mixture of open and closed states were only observed under turnover conditions (Kravchuk et al., 2022). This confirms that the closed state is a catalytically relevant active state and supports the hypothesis that open states are not solely a consequence of deactivation. However, to demonstrate that the open states are catalytically relevant other possible explanations for the observation of open states must be ruled out. Thus, it is important to note that another recent structural study of *E. coli* CI found that intact biochemically active preparations contained a significant fraction of broken particles on the cryoEM grids (Kolata and Efremov, 2021). After classification Kolata and Efremov found that most *E. coli* CI particles used for the reconstruction of the PA (151,357 vs. 134,976 particles) were particles that had completely dissociated from the MA (Kolata and Efremov, 2021). The biochemical preparations for the Kravchuk *et al*. and Kolata and Efremov studies are different and Kravchuk *et al*. do not report any disrupted particles (Kravchuk et al., 2022). Nonetheless, the results of Kolata and Efremov are a stark reminder that structural studies on extracted membrane protein complexes exist within a spectrum of biochemical stability and that complexes that are intact according to size exclusion chromatography, mass photometry and activity assays, can still end up as broken particles upon cryoEM grid preparation (Han et al., 2022; Kolata and Efremov, 2021). Therefore, it is not unreasonable to consider that the interaction with the air-water interface during grid preparation may act to convert closed state CI to the lower energy (higher entropy due to disorder and flexibility) open state.

Therefore, interactions with the air-water interface should be noted as an alternate explanation as to why most structures of CI across species appear to have open states. Notable exceptions so far are: 1) *Tetrahymena* CI, which is only found in a single closed resting state and has a much larger interaction interface between the PA and MA that stabilizes their association (Zhou et al., 2022); and 2) the thermophilic yeast *C. thermophilum* CI, which is found in multiple resting states, none of which corresponding to the open state seen in other species (Laube et al., 2022). In addition to thermophilic proteins being more stable in general, additional interactions between the PA and MA are also seen in *C. thermophilum* CI due to an N-terminal extension on NDUFA5. Although *Dm*-CI adopts an open-like state, it may be more resistant to disruption during grid preparation due to the expanded interaction between NADUFA5 and NDUFA10 (Figure 3B). However, each of these species need to be further characterized in the presence of substrates before general conclusions can be drawn.

In conclusion, the structure of *Dm*-CI provides the first structure of a protostomian CI that does not share the standard A-to-D transition as defined biochemically. Overall, with a few notable differences, the structure of *Dm*-CI is like that of mammals, validating its use as a genetically tractable model for the study of metazoan CI physiology. Given that inhibitors of CI have been developed as potential agricultural pesticides (Murai and Miyoshi, 2016), this structure will be a valuable resource for the development of more selective inhibitors. Due to its close relatedness to *D. melanogaster* (Figure 5-figure supplement 3), our structure is of particular value to develop targeted pesticides against spotted-wing drosophila (*D. suzukii*), a major invasive agricultural pest of the berry and wine industry in Southeast Asia, Europe and America (Tait et al., 2021). More broadly, the unique features of *Dm*-CI revealed here suggest strategies for the development of insecticides that could help control insect vectors of human disease. In general, this study highlights the utility of diverse model organisms in the study of important biochemical processes as we learn as much from the differences as we do from the similarities. In addition, our structures reveal unanticipated mechanisms that have evolved to regulate the assembly and activity of mitochondrial CI that may be exploited to modulate assembly or activity in other organisms. Additional studies on *Dm*-CI as well as other species are needed to fully understand the different mechanism which have evolved to regulate the assembly and activity of this important enzyme.

## Acknowledgments

The data were collected at the UC Davis BioEM Core facility. We thank Dr. Fei Guo for assistance with data collection. We thank Dr. María Maldonado for critical reading of the manuscript.

## Author contributions

Abhilash Padavannil, Methodology, Formal analysis, Investigation, Visualization, Validation, Writing - original draft, Writing - review and editing; Anjaneyulu Murari, Methodology; Shauna-Kay Rhooms, Methodology; Edward Owusu-Ansah, Conceptualization, Methodology, Writing - review and editing, Funding acquisition; James A. Letts, Conceptualization, Data curation, Formal analysis, Supervision, Funding acquisition, Validation, Visualization, Writing - original draft, Project administration, Writing - review and editing

## Data and materials availability

Single-particle cryogenic electron micrograph movies are available on the Electron Microscopy Public Image Archive, accession code 11272. The maps and models are available on the Electron Microscopy Database (EMDB) and Protein Data Bank (PDB). The accession codes for Locked open state are EMDB-28582, PDB-8ESZ and for the closed state are EMDB-28581, PDB-8ESW.

## Figure-figure supplements

**Video 1.** CryoEM density map and model of D. melanogaster CI. The subunits are colored as in Figure 1.

**Video 2**. CryoEM density map of *D. melanogaster* CI. Subunit NDUFA3 identified from the map is highlighted. The subunits are colored as in Figure 1.

**Video 3**. Structural analysis and comparison of NDUFS1-NDUFA2 interface in *Y. lipolytica* (PDB:6YJ4)*, D. melanogaster* (this study) and *O. aries* (PDB:6ZKC).

**Video 4.** Structural analysis of ND2, comparison of ND2-NDUFA10 interface and comparison of ND2-C2 interface in *D. melanogaster* (this study) and *O. aries* (PDB:6ZKC).

**Video 5.** Structural analysis of NDUFA11 subunit of *Dm*-CI and structural comparison of NDUFA11 in *Y. lipolytica* (PDB:6YJ4)*, D. melanogaster* (this study) and *O. aries* (PDB:6ZKC).

**Video 6.** 3D variability analysis of *D. melanogaster* CI, component 1. The 3DVA volumes are shown as a continuous movie. The movie emphases on the hinge region of PA/MA interface. Dm-CI subunits are colored as in Figure 1.

**Video 7.** 3D variability analysis of *D. melanogaster* CI, component 1. The 3DVA volumes are shown as a continuous movie. The movie emphases on the Q-site and interface loops at the PA/MA interface. *Dm*-CI subunits are colored as in Figure 1.

## Materials and methods

### *Drosophila* stocks and husbandry

*Drosophila* strains were maintained in vials containing agar, yeast, molasses and cornmeal medium supplemented with propionic acid and methylparaben in humidified environmental chambers (Forma environmental chambers) on a 12-h:12-h dark: light cycle. Mitochondrial preparations used for structure determination were from female *w^1118^*flies. To examine the extent of incorporation of NDUFA2-FLAG into CI, genetic crosses were set up between female flies of the genotype, *y w; Mhc-Gal4* and *UAS-NDUFA2-FLAG* males at 25°C. After the *Mhc-Gal4/UAS-NDUFA2-FLAG* flies eclosed, they were maintained at 25°C for one week, prior to dissection of thoraces. *Mhc-Gal4/ w^1118^* flies were used as controls.

### Mitochondria purification

Mitochondrial purification was performed as previously described (Rera et al., 2011). Briefly, fly thoraxes were dissected and gently crushed with a Dounce homogenizer in 1 mL per 20 thoraxes of pre-chilled mitochondrial isolation buffer containing 20 mM HEPES-KOH pH 7.5, 0.6 M Sorbitol, 1 mM EDTA, 1 mM DTT, 0.1 mg/ml BSA, 10 units/ml Trasylol and 0.5 mM PMSF on ice. After two rounds of centrifugation at 500 x g for 5 minutes at 4 °C to remove insoluble material, the supernatant was recovered and centrifuged at 5000 x g for 20 minutes at 4 °C. The pellet which is enriched for mitochondria was washed twice in the mitochondrial isolation buffer and stored at -80 °C until further processing.

### Spectroscopic assays for complex I’s active-to-deactive (A/D) transition

CI activity was measured by spectroscopic observation of the oxidation of NADH at 340 nm in 96-well plates at room temperature using a Molecular Devices (San Jose, CA) Spectramax M2 spectrophotometer. Measurements of the initial rates were done in 3-7 replicates, averaged and background corrected. An extinction coefficient of 6.22 mM^−1^ cm^−1^ (NADH) was used in the calculations. The reaction master mix consisted of 20 mM HEPES, pH 7.4, 50 mM NaCl, 10% glycerol (v/v), 0.1% BSA (w/v), 0.1% CHAPS (w/v), 0.1% digitonin (w/v), 200 μM DQ. *D. melanogaster* or porcine mitochondrial membrane samples were added to the corresponding mix at 50 μg/mL or 40 μg/mL respectively, mixed by tumbling and aliquoted into the 96-well plate to a total volume of 200 μl. For the NEM assay, the plate was incubated at 37°C for 10 min, after which 5 μM NADH or equivalent amount of buffer was added to the well and mixed by pipetting. 30 seconds after this addition, 0.5 mM NEM, 2 mM NEM or water was added to the corresponding wells and mixed by pipetting after which the plate was incubated for 20 min at room temperature and covered from light. The reactions were started by the addition of 300 μM NADH and briefly mixed by pipetting before recording every 2 s for 5 min.

### Western blotting

Western blotting was performed as previously described (Murari et al., 2022). Briefly, following the separation of protein complexes on 3–12% precast Bis-Tris Native PAGE gels (Life Technologies), the proteins were transferred to polyvinylidene difluoride (PVDF) membranes (Bio-Rad). Subsequently, the PVDF membrane was blocked in 5% (wt/vol) nonfat dry milk (NFDM) in tris-buffered saline (TBS) for 30 min and incubated in the appropriate primary antibody dissolved in 2% BSA and 0.1% Tween 20 in TBS (TBST) overnight at 4°C. Subsequently, the blot was rinsed four times for 10 min each in 0.1% TBST, blocked for 30 min in 5% (wt/vol) NFDM in TBST, and incubated for 2 h at room temperature with the appropriate HRP-conjugated secondary antibody dissolved in 2% BSA and 0.1% TBST. Afterwards, samples were rinsed four times for 10 min each in 0.1% TBST. Immunoreactivity was detected by a SuperSignal West Pico PLUS Chemiluminescent kit (Thermo Scientific, 34578) and analyzed by a ChemiDoc gel imaging system from Bio-Rad. The primary antibodies used were anti-NDUFS3 (Abcam, ab14711), anti-NDUFA2 (this study), anti-FLAG (MilliporeSigma, F3165) and anti-ATPsynß (Life Technologies, A21351). Secondary antibodies used were goat anti-rabbit horseradish peroxidase (PI31460 from Pierce) and goat anti-mouse horseradish peroxidase (PI31430 from Pierce). To generate a rabbit polyclonal antibody for the *Drosophila* ortholog of NDUFA2 (CG15434) the following synthetic peptide was used: DPKGDTSKGVREYVER-Cys.

### Electron transport chain complex purification

The following operations were carried out at 4°C unless otherwise indicated. The mitochondria pellet was resuspended and lysed in milli-Q water at 10 mL/g (of starting mitochondria, wet weight) using a Dounce homogenizer, to which KCl was added to a final concentration of 150 mM. The mitochondrial membrane was pelleted by centrifugation at 32,000 x g for 45 minutes and washed once in buffer M10 (20 mM Tris pH 7.4, 50 mM NaCl, 1 mM EDTA, 2 mM dithiothreitol (DTT), 0.002% PMSF (w/v), 10% glycerol (v/v)) at 18 mL/g (of starting mitochondria). The resulting membrane pellet was resuspended in buffer M10 at 3 mL/g (of starting mitochondria) and the protein concentration was determined using a BCA assay (Pierce Thermo Fisher). The resuspended membranes were stored at 10 mg/mL of total protein at -80°C in a final glycerol concentration of 30% (v/v) after dilution with buffer M90 (20 mM Tris pH 7.4, 50 mM NaCl, 1 mM EDTA, 2 mM DTT, 0.002% PMSF (w/v), 90% glycerol (v/v)). Usual yield was ∼30 mg total membrane protein per gram of *D. melanogaster* thorax.

The thawed mitochondrial membrane resuspension was solubilized in buffer MX (30 mM HEPES pH 7.7, 150 mM potassium acetate, 0.002% PMSF, 10% (v/v) glycerol) by slow tumbling for 1 hour at 4°C with 1% digitonin (w/v) at a detergent-to-protein ratio of 4:1 (w/w). The insoluble material was cleared by centrifugation at 16,000 x g for 20 minutes. Amphipol A8-35 was added to the supernatant to a final concentration of 0.3% (w/v), before incubation with slow tumbling for 1 hour at 4°C. Digitonin was removed from the supernatant by dialyzing the sample first in a buffer containing γ-cyclodextrin followed by dialysis in buffer containing Bio-beads. The dialyzed sample was centrifuged at 16,000 x g for 20 minutes to remove any precipitate. The supernatant was concentrated in 100 kDa MWCO centrifugal concentrators to 0.250 mL and loaded on to a continuous 15% to 45% (w/v) sucrose gradient in SGB buffer (15 mM HEPES pH 7.8, 20 mM KCl). After centrifugation in a SW40Ti swinging-bucket rotor at 149,176 x g for 24 hours, the sucrose gradients were fractionated using a Biocomp gradient profiler. Fractions were assayed for CI activity by running them on a 3%-12% Tris-glycine blue-native PAGE (BN-PAGE) gel and a nitrotetrazoleum blue in-gel assay was performed as previously described (Maldonado et al., 2020). Fractions displaying CI activity were pooled and concentrated to a final concentration of 5 mg/mL.

### CryoEM grid preparation and data collection

Four microliters of concentrated fractions from the sucrose gradients were applied onto a Quantifoil R1.2/1.3 300 mesh copper grids glow-discharged at 30 mA for 30 seconds before sample application. In a GP2, the grid was first incubated for 20 seconds at 100% humidity, then blotted for 4 seconds before plunge-freezing into liquid ethane cooled by liquid nitrogen. A total of 11,065 movies were collected using SerialEM on a 200 kV ThermoFisher Glacios microscope equipped with a Gatan Quantum K3 detector, at a nominal magnification of 56,818 (0.44 Å/pixel under super-resolution mode). A dose of 20 electrons/Å^2^/s with 3-second exposure time was fractionated into 75 frames for each movie.

### CryoEM image processing

The raw movies were binned 2-fold and motion-corrected using the MotionCor2 (Zheng et al., 2017), followed by per-micrograph contrast transfer function (ctf) estimation using the CTFFIND4.1 (Rohou and Grigorieff, 2015), both implemented in Relion 3.1.0 (Zivanov et al., 2018a). Micrographs were then curated to remove images lacking high resolution ctf correlations. Particles were picked using crYOLO (Wagner et al., 2019). The initial 698,452 picked particles were extracted in Relion 3.1.0 with 512 pixel^2^ boxes, followed by 2D classification, 3D *ab initio* reconstruction and 3D refinement in cryoSPARC v3.2.0 (Punjani et al., 2017). Iterative 2D classification and 3D *ab initio* reconstruction resulted in 293,389 good particles corresponding to Dm-CI, 25,080 particles corresponding to Dm-CIII and 31,198 particles corresponding to Dm-CV (Fig 1-figure supplement 2). Homogenous refinement followed by non-uniform refinement (Punjani et al., 2020) of *Dm*-CI in cryoSPARC resulted in an initial reference map of 3.71 Å. This particle set was then transferred back into Relion 3.1.0 for further processing involving several rounds of global search, CTF refinement, Bayesian polishing (Zivanov et al., 2018b) and local searches resulting in a final map of 3.44 Å. This map was used for initial model building in Coot (Emsley et al., 2010). Following initial model building and refinement in Phenix (Liebschner et al., 2019), masks corresponding to the peripheral arm, membrane arm and the whole CI were generated in Relion. Iterative masked refinement and 3D classification resulted in a final reference map of 3.30 Å of the *Dm*-CI after import back into cryoSPARC for non-uniform refinement.

Poor local resolution and broken density around at the matrix interface of the MA and PA (NDUFA10, NDUFA5, NDUFA6, NDUFS4, NDUFAB1-α) of CI prompted us to further classify CI particles using mask around the hinge region. 3D classification of CI particles using mask around the hinge region resulted in two distinct classes of CI particles. Iterative homogenous refinement and non-uniform refinement of the classes resulted in reference maps of 3.40 Å for both classes. The final focused map was post-processed using DeepEMhancer (Sanchez-Garcia et al., 2021) which improved the connectivity of certain regions of protein but also removed density for structured lipids. All software suites used for data processing and refinement except for cryoSPARC were accessed through the SBGrid consortium (Morin et al., 2013). 3D variability analysis (3DVA) on all 239,389 good particles was performed in cryoSPARC (Punjani and Fleet, 2020) to solve for three eigen volumes of the 3D covariance. Volume series corresponding to each of the components is generated in cryoSPARC. Molecular graphics and analyses were performed with UCSF ChimeraX, developed by the Resource for Biocomputing, Visualization, and Informatics at the University of California, San Francisco, with support from National Institutes of Health R01-GM129325 and the Office of Cyber Infrastructure and Computational Biology, National Institute of Allergy and Infectious Diseases (Pettersen et al., 2021).

### Model building and refinement

All manual model building was performed in Coot 0.9.2 (Emsley et al., 2010) and refinements were performed in Phenix-1.19.1 (Liebschner et al., 2019). Mammalian CI was docked into the *Dm*-CI map and Alpha-fold models (Jumper et al., 2021), accessed via Uniprot (Consortium et al., 2020), of *Dm*-CI subunits were structurally aligned to each of the corresponding mammalian CI subunits to generate an initial model of the *Dm*-CI. The model-map fit was manually inspected, and the model was rebuilt where necessary to generate an initial *Dm*-CI model. Secondary structure restraints were first automatically generated from the manually built model, then edited according to the outcome of the Phenix refinement. Bond length and angle restraints for metal ion coordination and amino acid side chain linkage were generated manually, and a ligand.cif file was also provided for non-default ligands in Phenix. The refined model was manually inspected and edited in Coot before the next round of Phenix refinement, and this iterative cycle continued until the model statistics converged before submission of maps and models to the EMDB and PDB databases. Model statistics and details by subunit are provided in Table 1.

---

**Figure 1-figure supplement 1.**
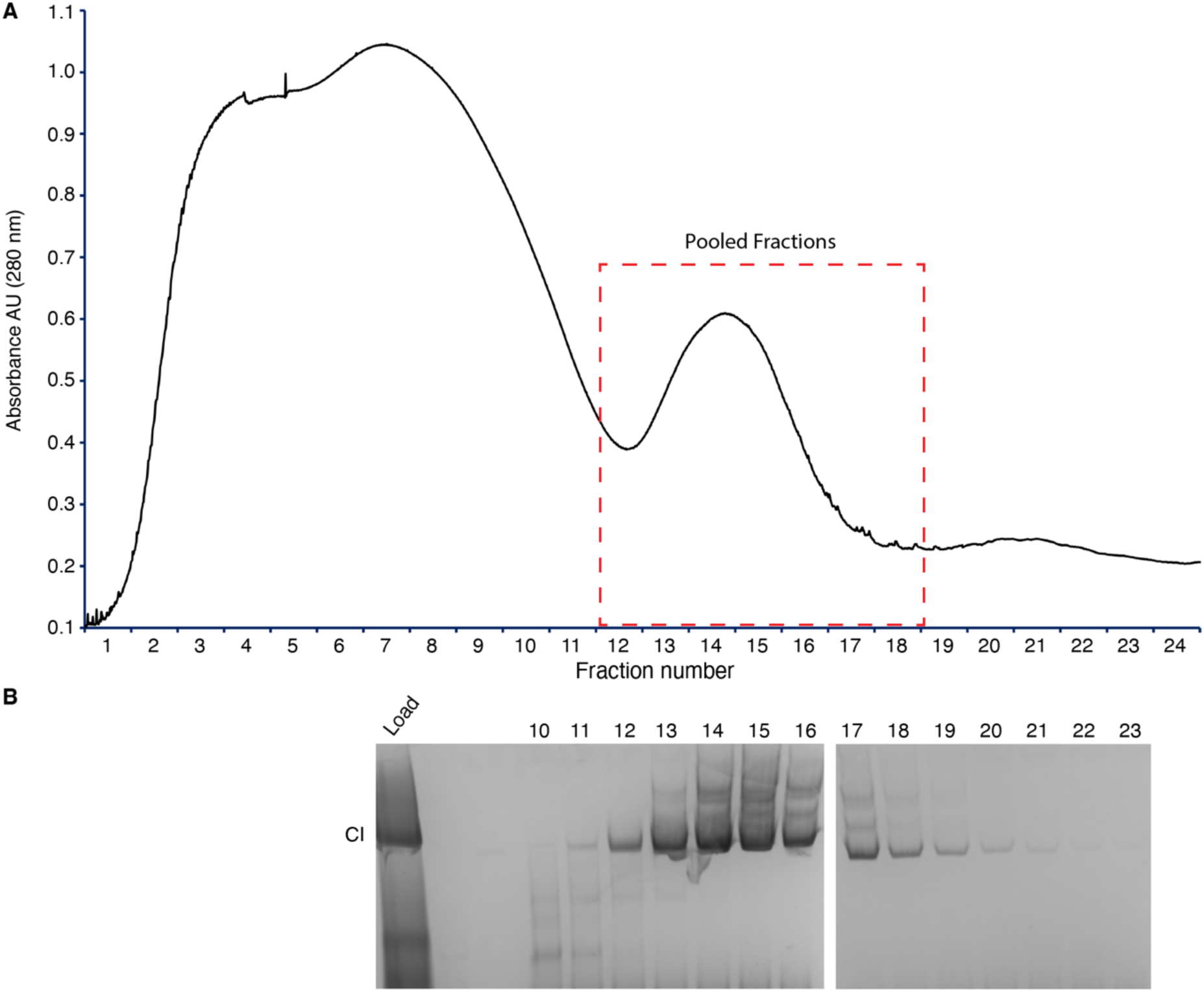
(A) Chromatogram of sucrose gradient fractionation of digitonin-extracted and amphopol A8-35-exchanged *D. melanogaster* mitochondrial complexes. (B) Blue-native PAGE (BN-PAGE) of fractions from (A) visualized by CI in gel activity staining.

**Figure 1-figure supplement 2.**
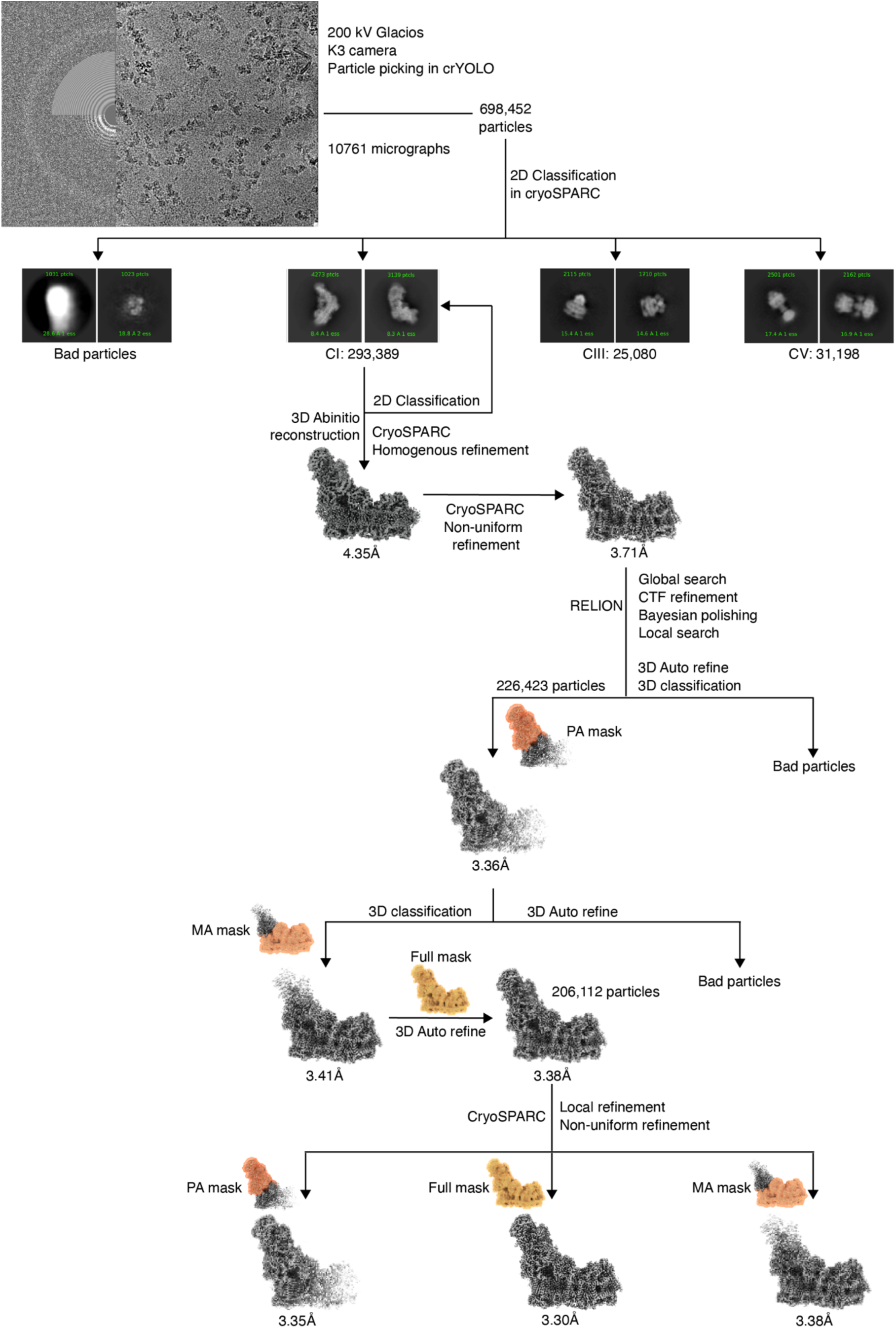
CryoEM image processing and refinement. A total of 10,761, movies were collected on a 200 kV Glacios microscope with K3 detector, from which 698,452 particles were initially picked using CrYOLO (Wagner et al., 2019). In CryoSPARC (Punjani et al., 2017), 2D classification revealed classes corresponding to CI, CIII2 and CV. After further particle classification via 3D *ab initio* reconstruction, a total of 293,389 *Dm*-CI particles were obtained. Homogenous refinement followed by non-uniform refinement (Punjani et al., 2020) resulted in an initial map of 3.71 Å. Further refinement with global and local CTF corrections and Bayesian polishing in RELION (Zivanov et al., 2020, 2019) followed by local refinement using masks around the *Dm*-CI peripheral arm, membrane arm and full mask in CryoSPARC, resulted in a final map of 3.30 Å.

**Figure 1-figure supplement 3.**
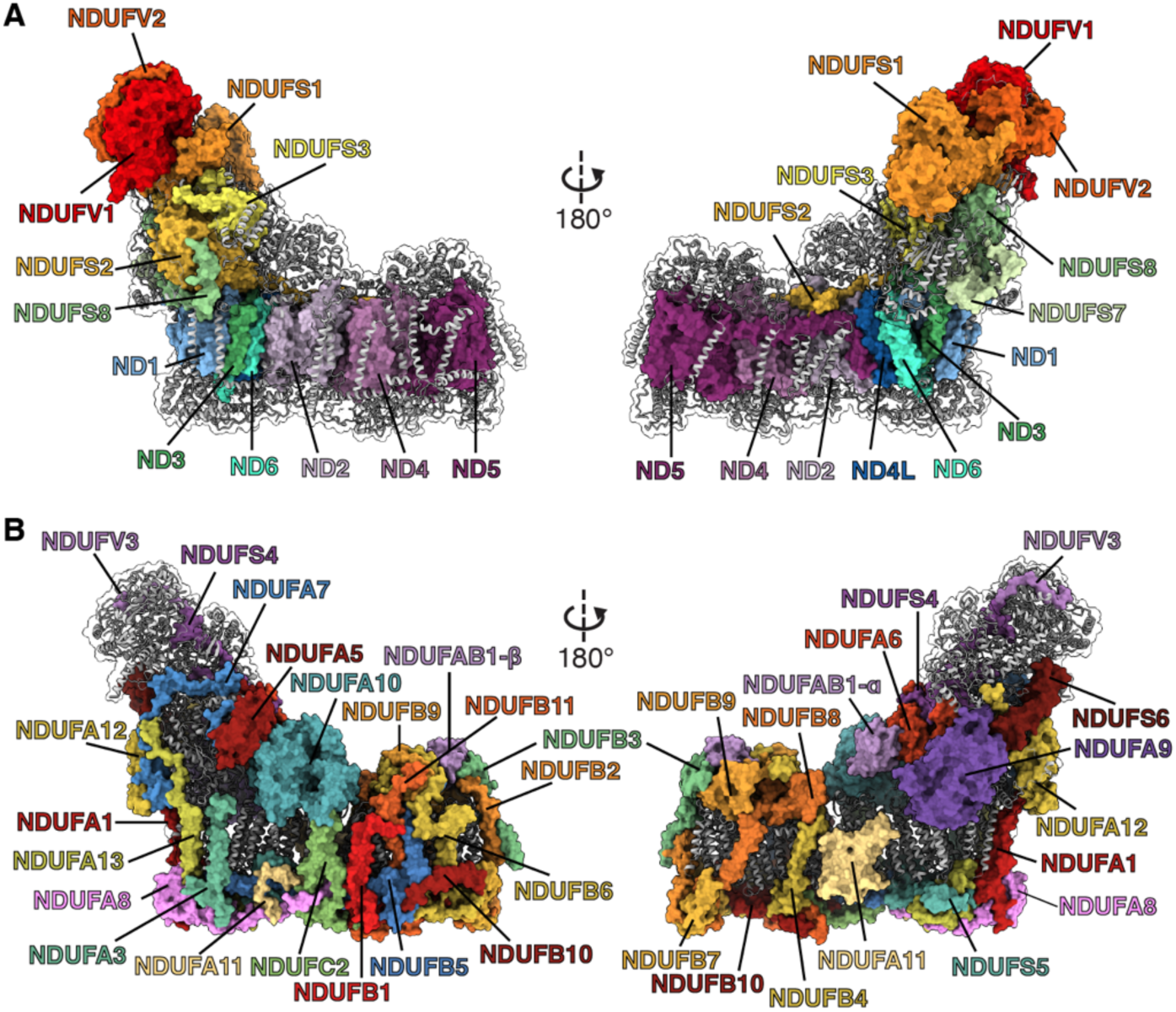
Overall structure of *Dm*-CI. (A) *Dm*-CI’s 14 core subunits in colored surface; accessory subunits are shown as grey cartoons. (B) *Dm*-CI’s 29 accessory subunits in colored surface; 14 core subunits are shown as grey cartoon. The subunits are colored as in Figure 1.

**Figure 1-figure supplement 4.**
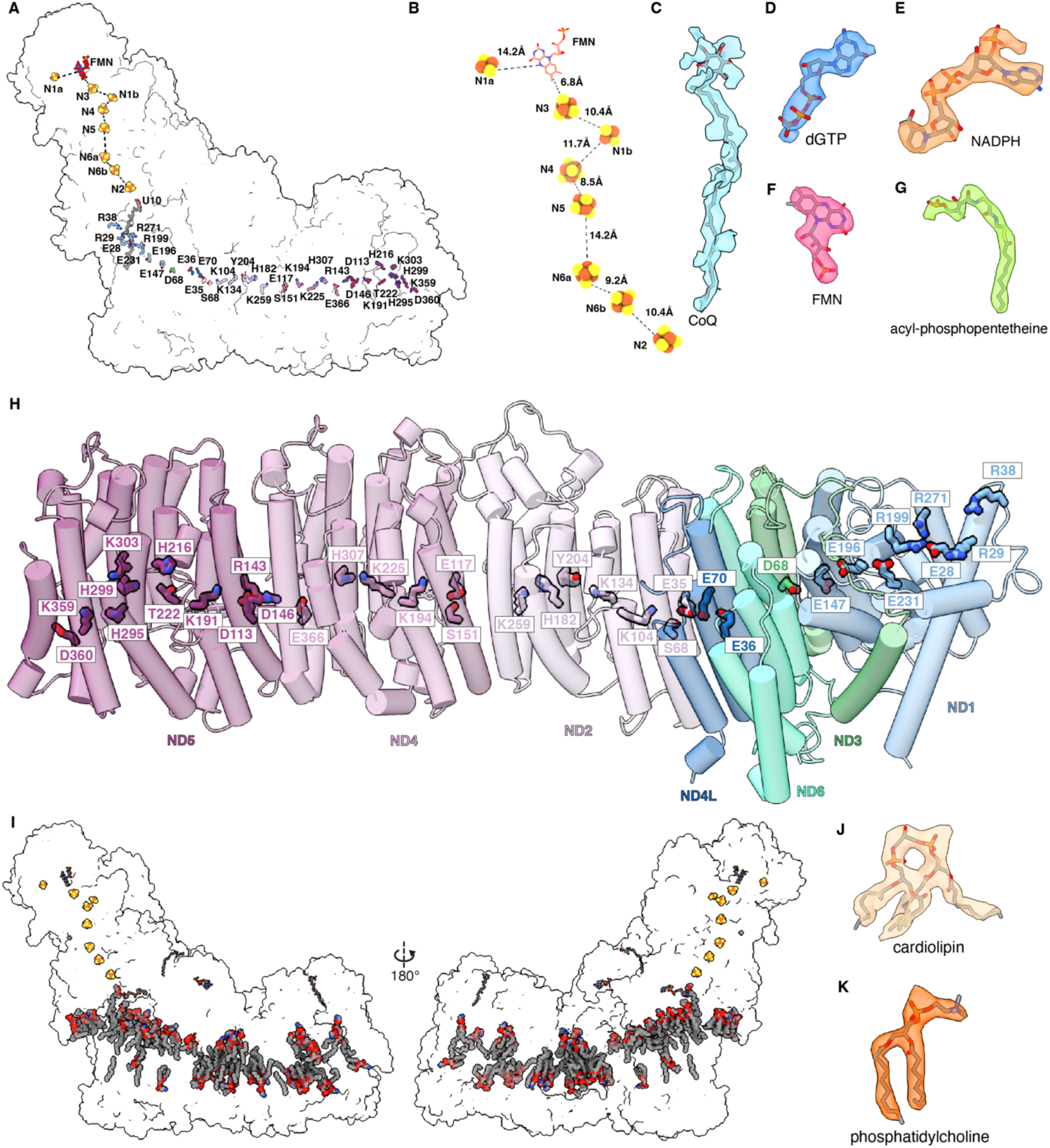
*Dm*-CI electron transfer pathway, co-factors and hydrophilic axis. (A) The electron transfer pathway and the conserved hydrophilic axis are shown in a transparent surface of the overall structure. (B) The redox cofactors of the electron transfer pathway of *Dm*-CI are shown. The FES clusters are shown as spheres colored by the element. The dashed lines indicate the edge-to-edge distances. (C D E F G) Density of *Dm*-CI co-factors: ubiquinone (CoQ), 2′-deoxyguanosine-5′-triphosphate (dGTP) in NDUFA10, NADPH in NDUFA9, flavin mononucleotide (FMN) in NDUFV1, and acyl-phosphopantetheine in NDUFAB1-α (H) The hydrophilic axis along the membrane arm of *Dm*-CI is show. Core subunits that form the membrane arm are shown as cylinders, colored as in Figure 1. The conserved residues that constitute the hydrophilic axis are shown in stick colored by subunit. (I) The distribution of lipids is shown. (J K) Representative density of lipids cardiolipin (CDL) (J) 1,2-diacyl-SN-glycero-3-phosphocholine (PC1) (K).

**Figure 1-figure supplement 5.**
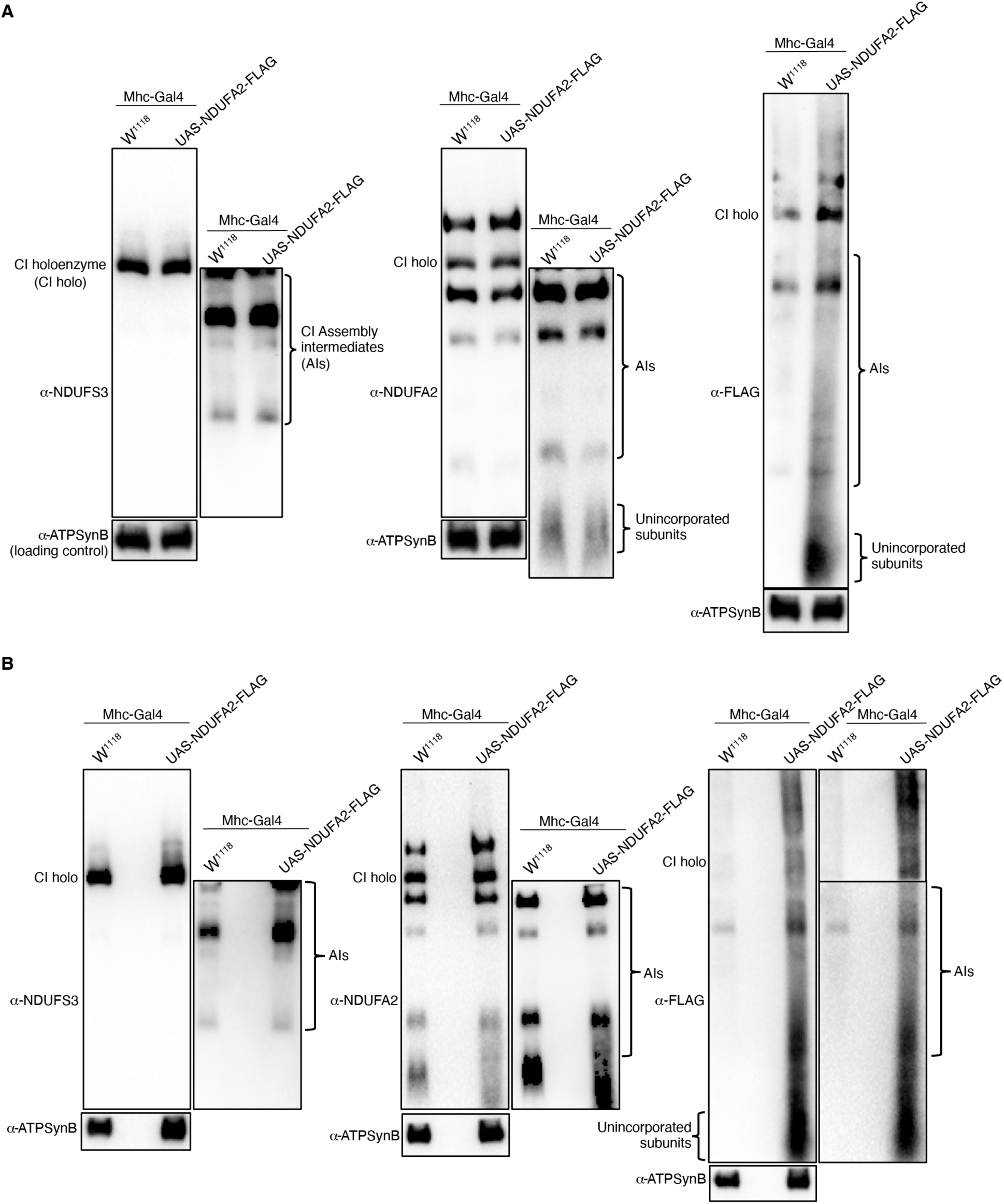
Mitochondrial preparations from thoraces of 1-week old flies with the genotypes indicated were analyzed by BN-PAGE, followed by Western blotting with the antibodies shown. Panels A and B are biological replicates.

**Figure 1-figure supplement 6.**
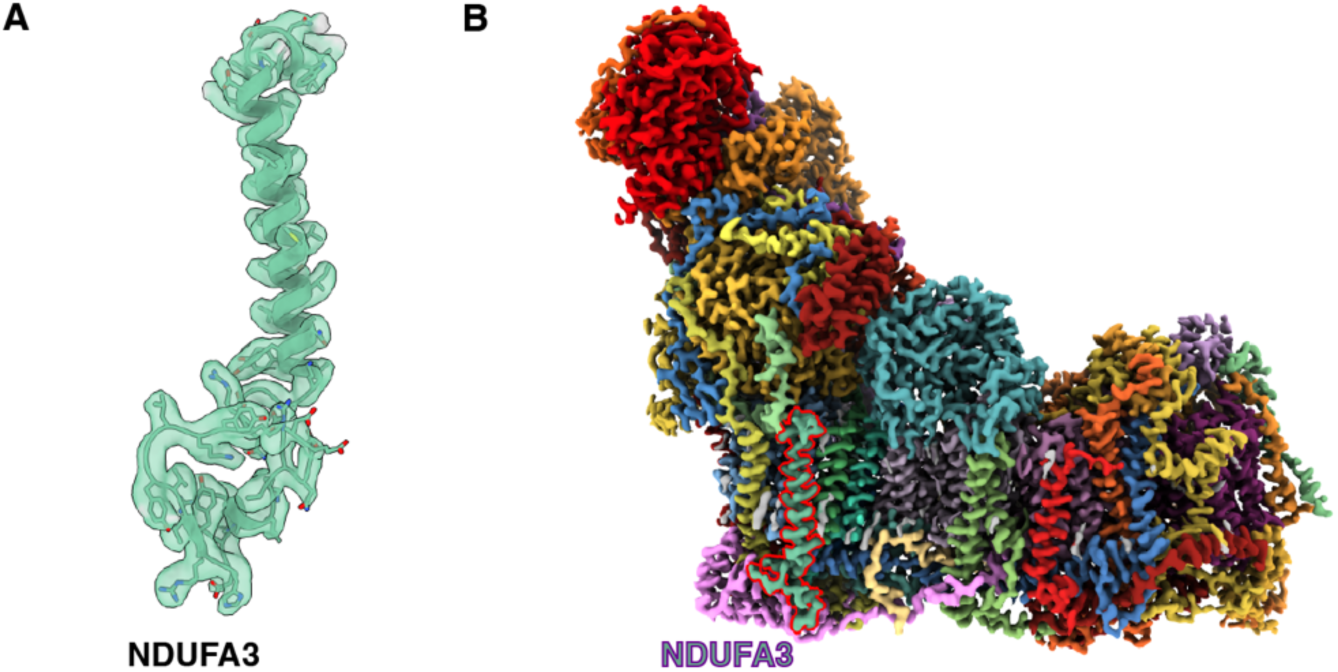
NDUFA3 of *Dm*-CI. (A) Newly identified NDUFA3 of *Dm*-CI is shown as cartoon embedded in the colored density. (B) The position of NDUFA3 density as identified in the overall *Dm*-CI is highlighted in red. The Cryo-EM map is colored according to the subunits as in Figure 1.

**Figure 1-figure supplement 7.**
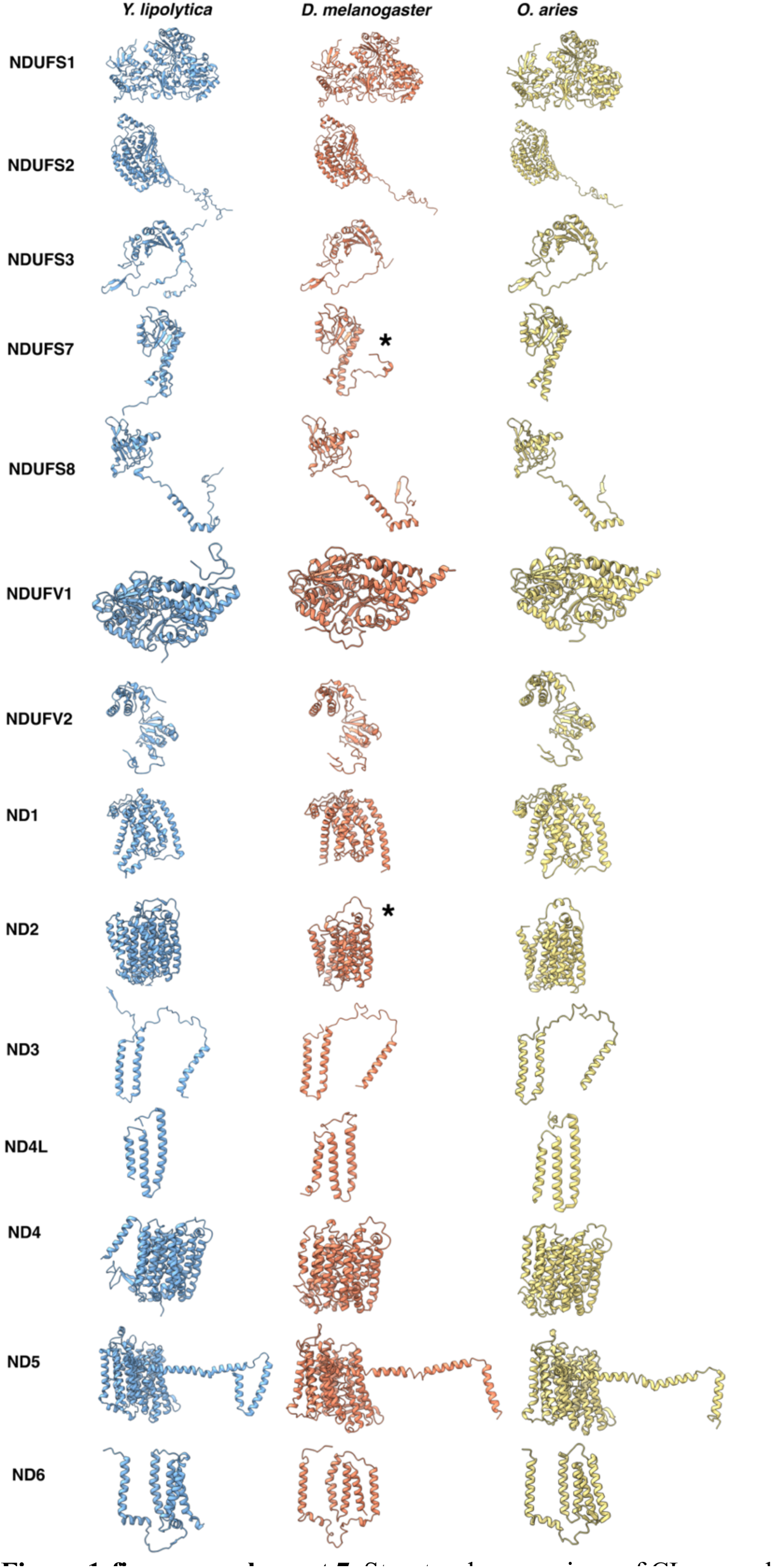
Structural comparison of CI core subunits from *Y. lipolytica* (PDB: 6RFR) *D. melanogaster* (this study) and *O. aries* (PDB:6ZKC). The subunits are shown as colored cartoons *Y. lipolytica* (blue) *D. melanogaster* (orange) and *O. aries* (yellow).

**Figure 2-figure supplement 1.**
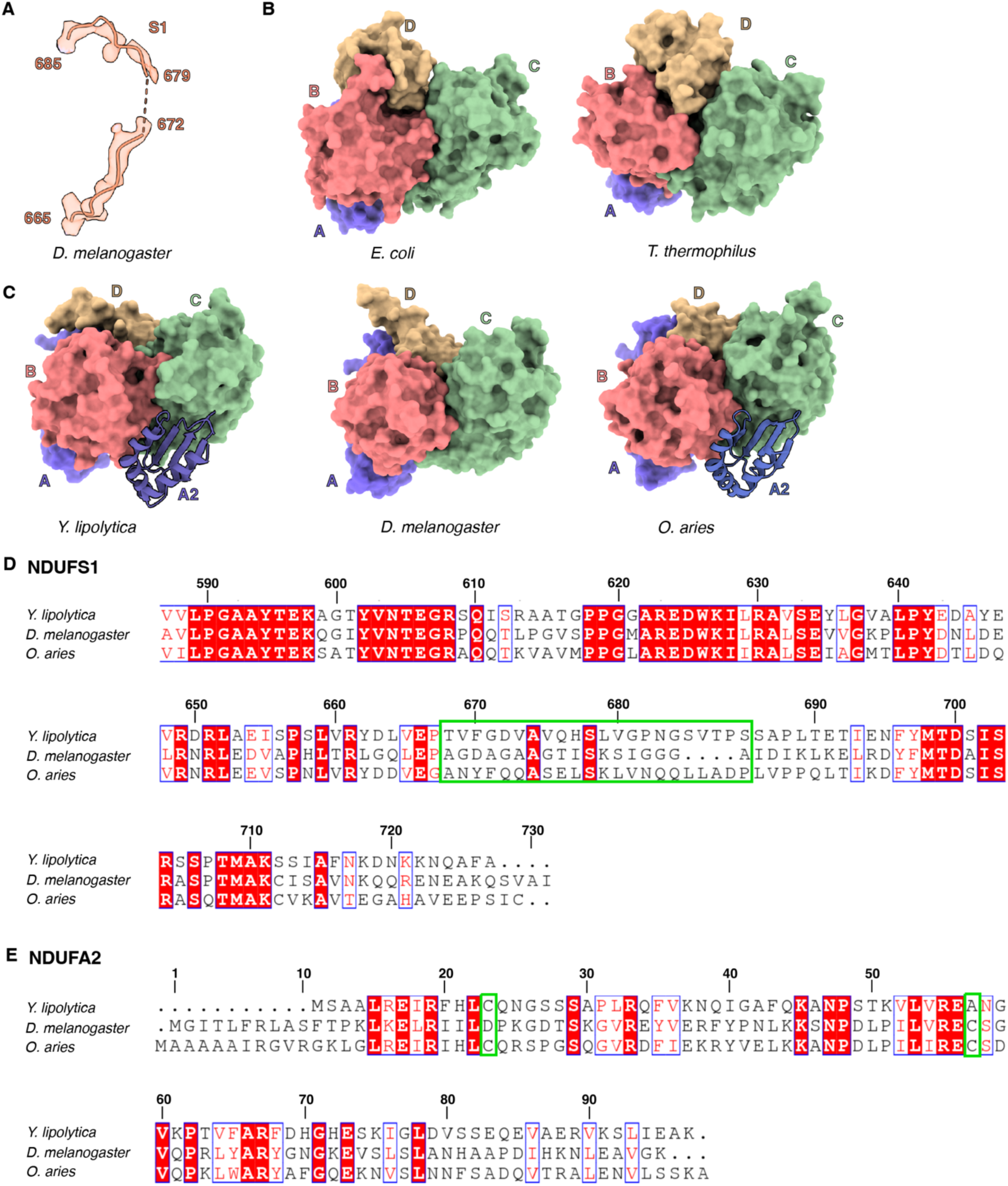
Structural analysis of NDUFS1-NDUFA2 interface. (A) NDUFS1 region 665-685 of *Dm*-CI is shown as cartoon embedded in the density (B) Domain architecture of NDUFS1 in *E. coli* (PDB: 7NZ1) and *T. thermophilus* (PDB: 4HEA) (C) Domain architecture of NDUFS1 in *Y. lipolytica* (PDB: 6YJ4), *D. melanogaster* (this study), and *O. aries* (PDB: 6ZKC). (D) Sequence alignment of C-terminal region of NDUFS1 from *Y. lipolytica, D*. *melanogaster* and *O. aries*. Numbered according to *Dm*-S1. The unstructured region of *Dm*-S1 is boxed green. (E) Sequence alignment of NDUFA2 from *Y. lipolytica, D. melanogaster* and *O. aries*. Numbered according to *Dm*-A2. Cysteines forming the di-sulfide bonds in oxidized form of NDUFA2 are boxed green.

**Figure 2-figure supplement 2.**
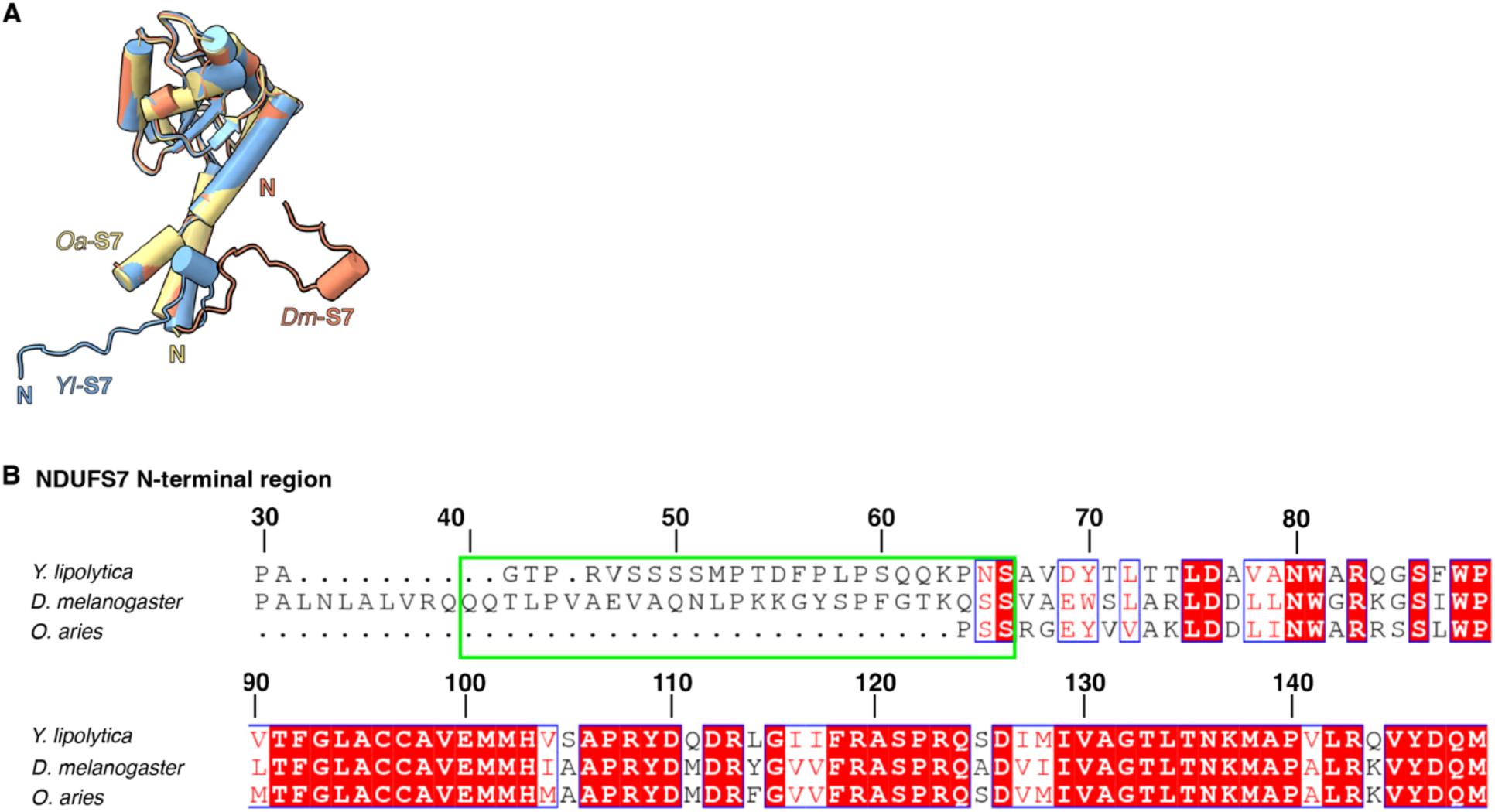
Structural analysis of NDUFS7. (A) Structural alignment of NDUFS7 from *Y. lipolytica* (PDB: 6YJ4), *D. melanogaster* (this study) and *O. aries* (PDB: 6ZKC) is shown. (B) Sequence alignment of N-terminal region of NDUFS7 from *Y. lipolytica*, *D. melanogaster* and *O. aries*. Numbered according to *Dm*-NDUFS7. The sequence in the N terminal extension is boxed green.

**Figure 2-figure supplement 3.**
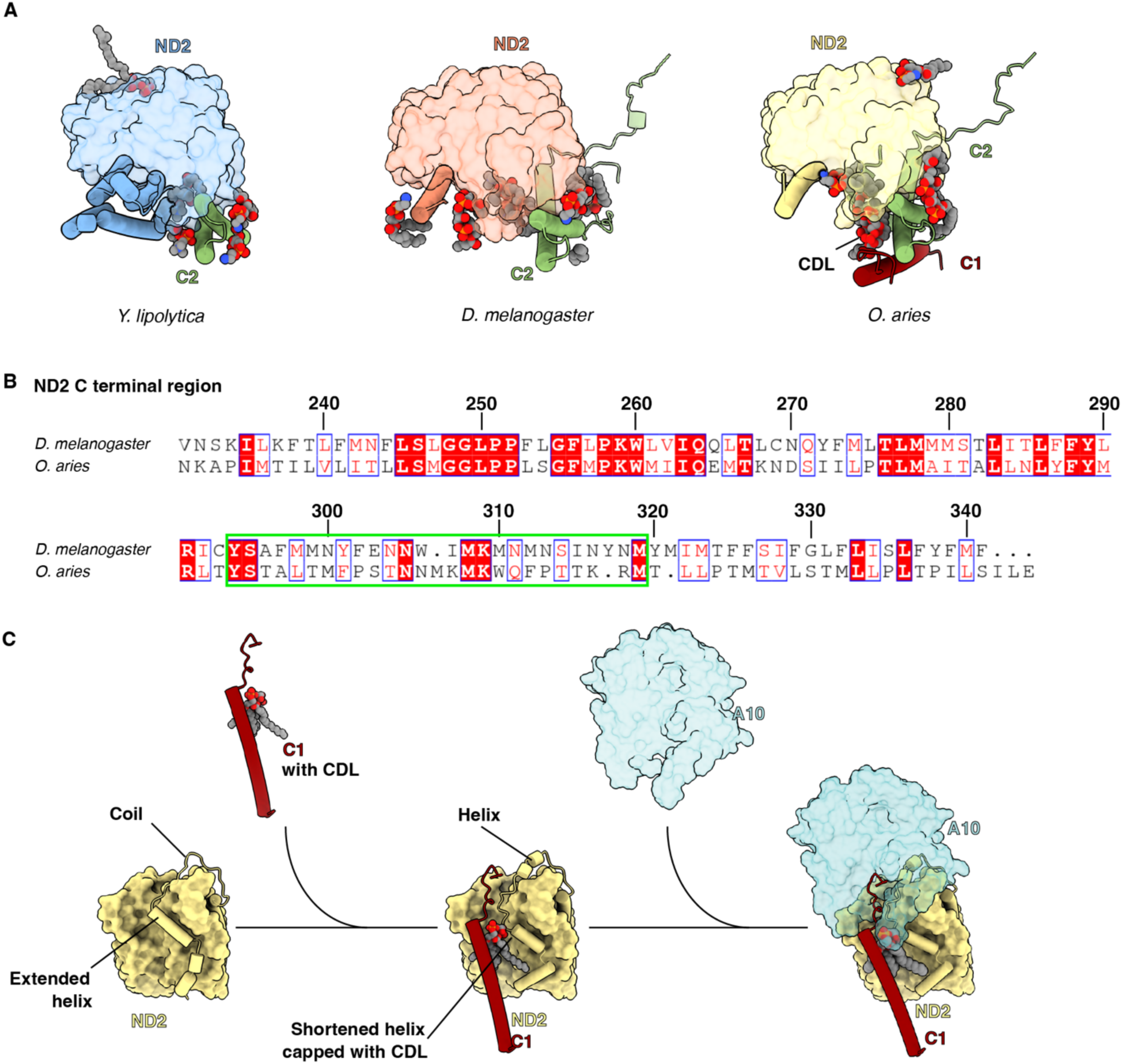
Structural analysis of ND2-NDUFA10 interface. (A) *Y. lipolytic* (PDB: 6YJ4), *D. melanogaster* (this study) and *O. aries* (PDB :6ZKC) ND2 is shown in surface. The N-terminal helices of ND2 are show as cartoons. NDUFC2 and NDUFC1 are shown as cartoons. Lipids are shown as spheres colored by element. (B) Sequence alignment of C-terminal region of ND2 from *D*. *melanogaster* and *O. aries*. Numbered according to *Dm*-ND2. The sequence of TMH10-11 loop region is boxed green. (C) Schematic representation of the role of NDUFC1 in mammalian CI assembly. TMH11^ND2^ has an extended helix. NDUFC1 brings in a cardiolipin to cap the shortened TMH11^ND2^, inducing the formation of short helices in the erstwhile coiled TMH10-11^ND2^ loop. The short helices form the interface for NDUFA10 to bind to complex I. NDUFC2 is not shown for clarity.

**Figure 2-figure supplement 4.**
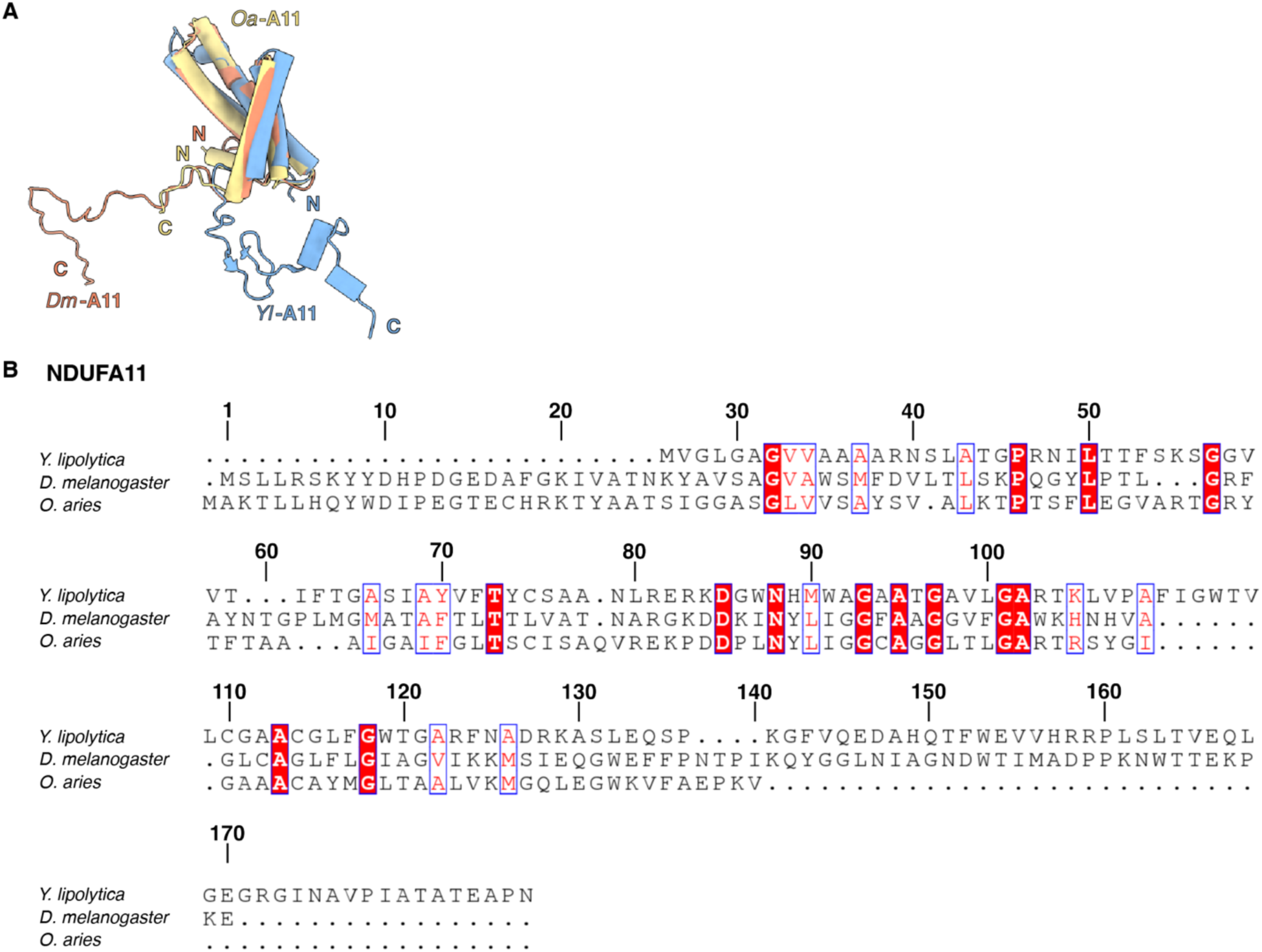
NDUFA11. (A) Structural alignment of NDUFA11 from *Y. lipolytica* (PDB: 6YJ4) *D. melanogaster* (this study) and *O. aries* (PDB: 6ZKC) is shown (B) Sequence alignment of NDUFA11 from *Y. lipolytica*, *D. melanogaster* and *O. aries*. Numbered according to *Dm*-NDUFA11.

**Figure 3-figure supplement 1.**
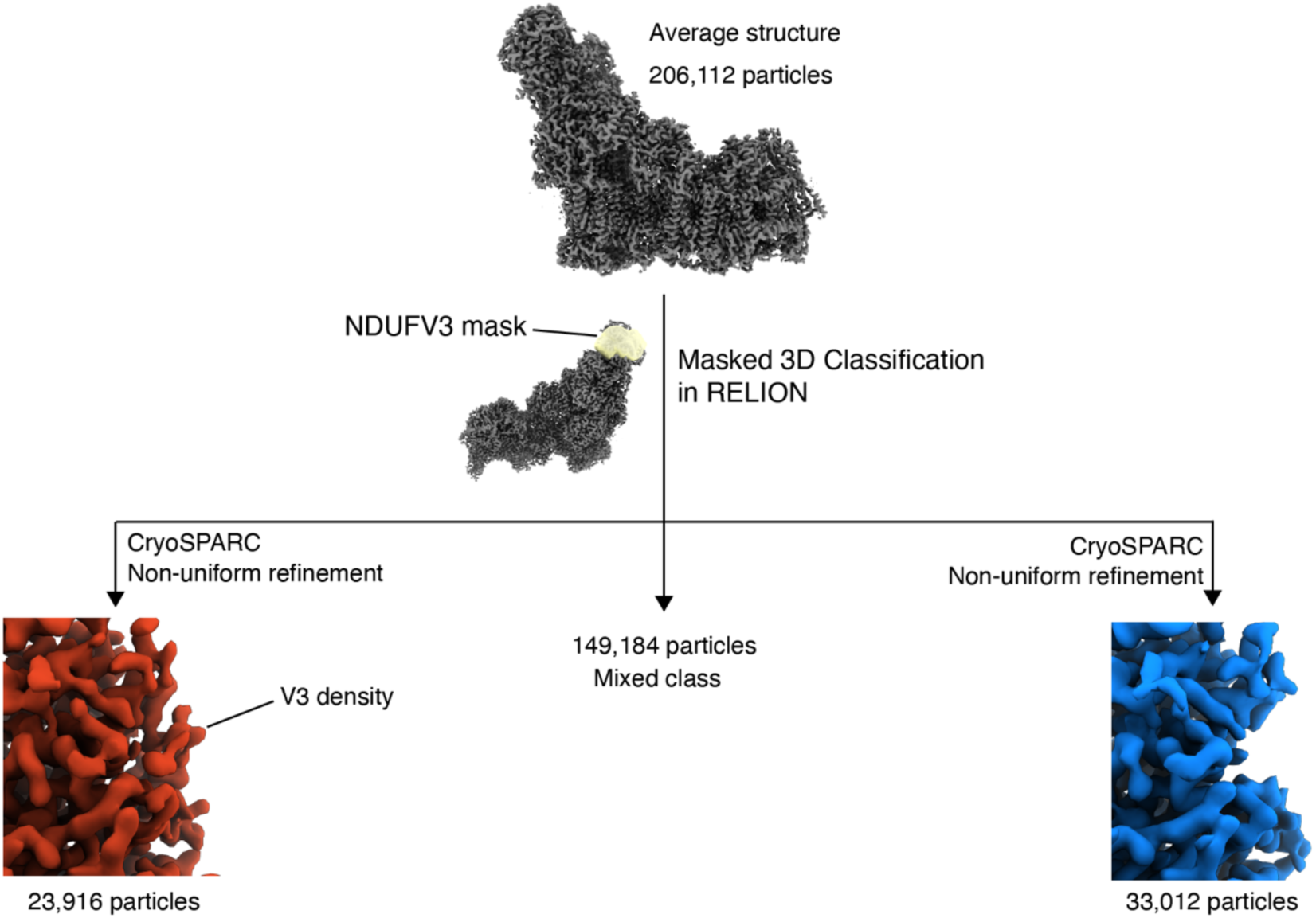
NDUFV3 is sub-stoichiometric. Masked classification of 206,112 *Dm*-CI particles using mask around the tip of the peripheral arm resulted in three distinct classes of particles, Class I with clear NDUFV3 density, Class II a set of mixed particles, Class III with clear absence of NDUFV3 density.

**Figure 3-figure supplement 2.**
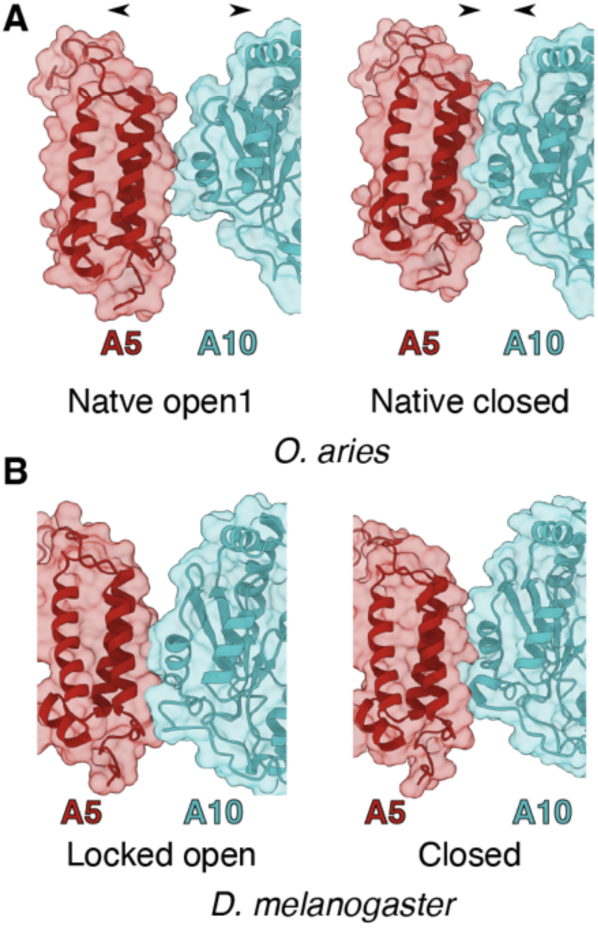
State dependent interactions between NDUFA5 and NDUFA10 (A) Transparent surface of open D-state (PDB:6ZKD) and closed A-state (PDB:6ZKC) of *Oa*-CI showing NDUFA10 and NDUFA5 as cartoons. (B) Transparent surface of Locked open and closed state of *Dm*-CI showing NDUFA10 and NDUFA5 as cartoons (this study).

**Figure 3-figure supplement 3.**
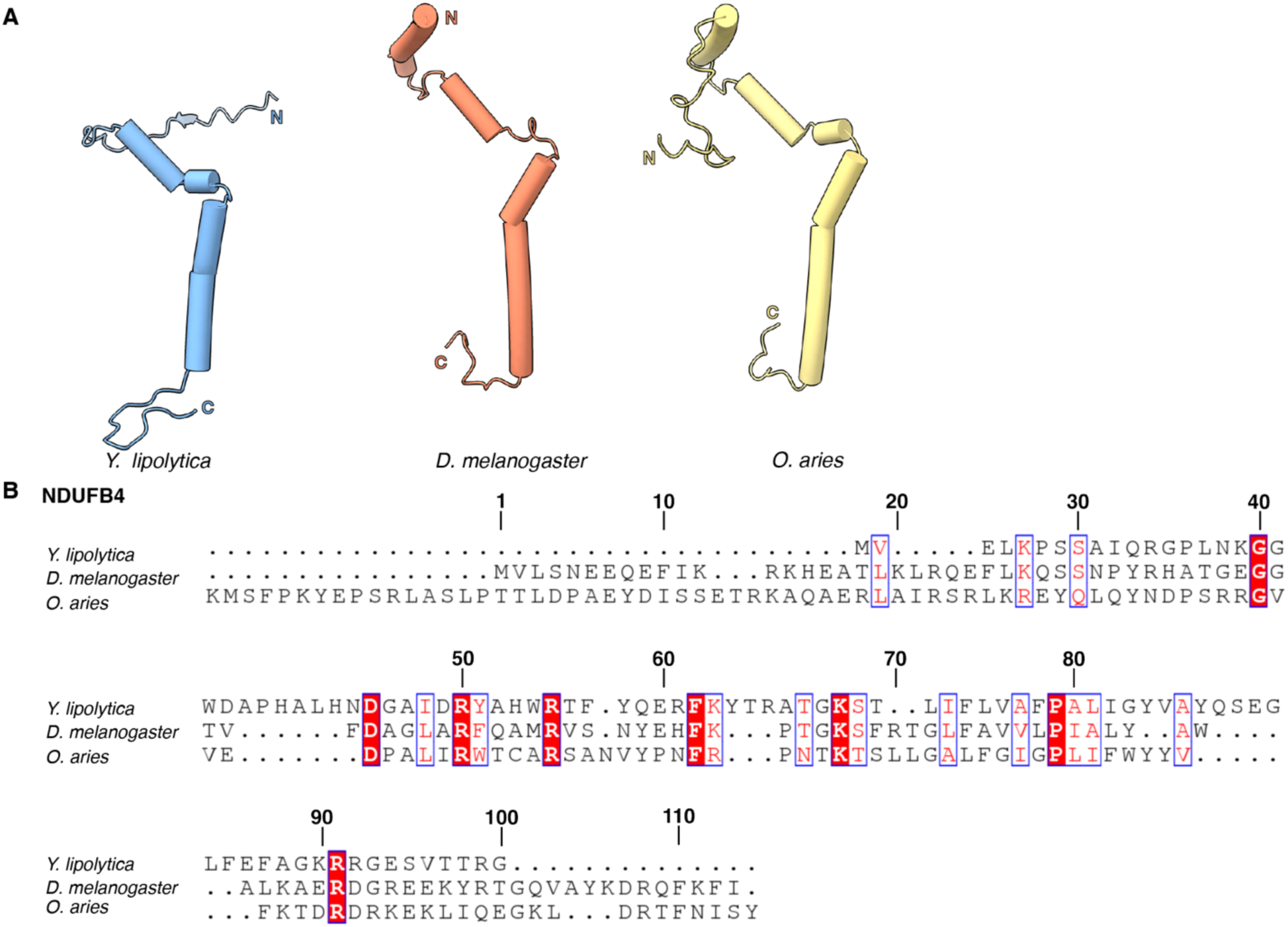
NDUFB4. (A) Structural comparison of NDUFAB4 from *Y. lipolytica* (PDB: 6YJ4) *D. melanogaster* (this study) and *O. aries* (PDB: 6ZKC) is shown (B) Sequence alignment of NDUFB4 from *Y. lipolytica*, *D. melanogaster* and *O. aries*. Numbered according to Dm-NDUFB4.

**Figure 3-figure supplement 4.**
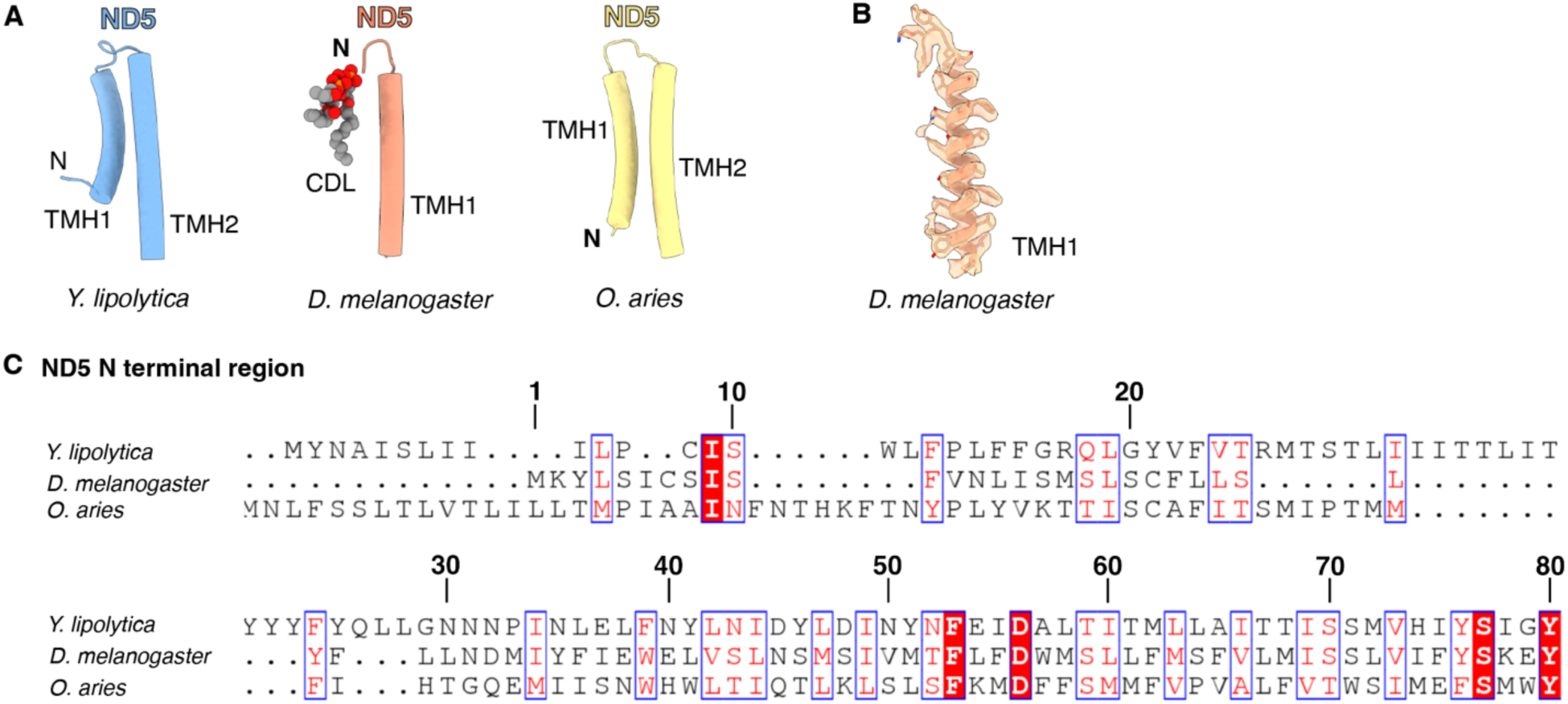
*Dm*-ND5 has a truncated N-terminus. (A) Structural comparison of N-terminal helices of ND5 in *Y. lipolytica* (PDB: 6YJ4) *D. melanogaster* (this study) and *O. aries* (PDB: 6ZKC). (B) *Dm*-ND5 TMH1 is shown as cartoon embedded in density map. (C) Sequence alignment of N-terminal region of ND5 from *Y. lipolytica, D. melanogaster*, and *O. aries*. Numbered according to *Dm*-ND5.

**Figure 3-figure supplement 5.**
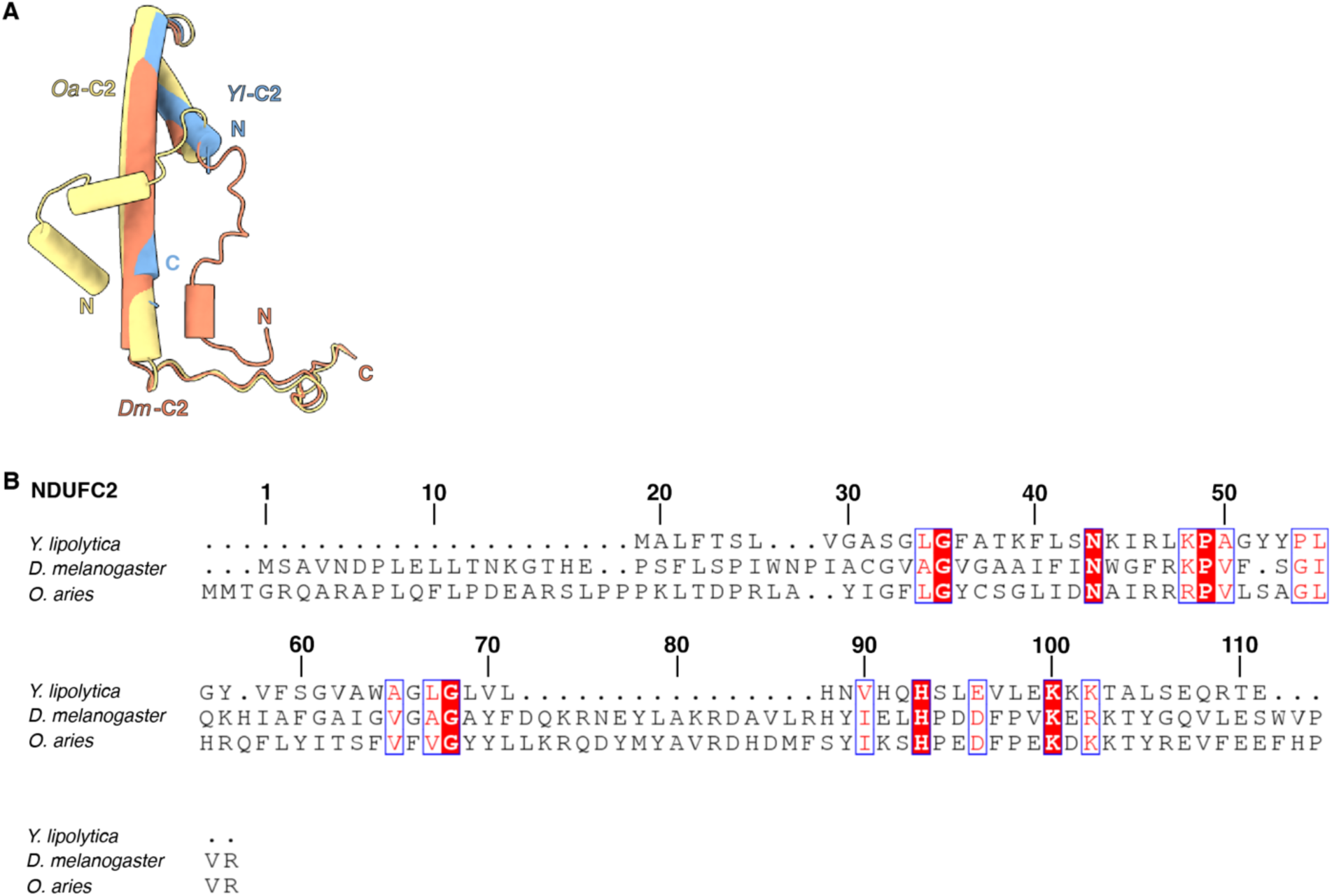
NDUFC2. (A) Structural alignment of NDUFC2 from *Y. lipolytica* (PDB: 6YJ4) *D. melanogaster* (this study) and *O. aries* (PDB: 6ZKC) is shown (B) Sequence alignment of NDUFC2 from *Y. lipolytica*, *D. melanogaster* and *O. aries*. Numbered according to *Dm*-NDUFC2.

**Figure 4-figure supplement 1.**
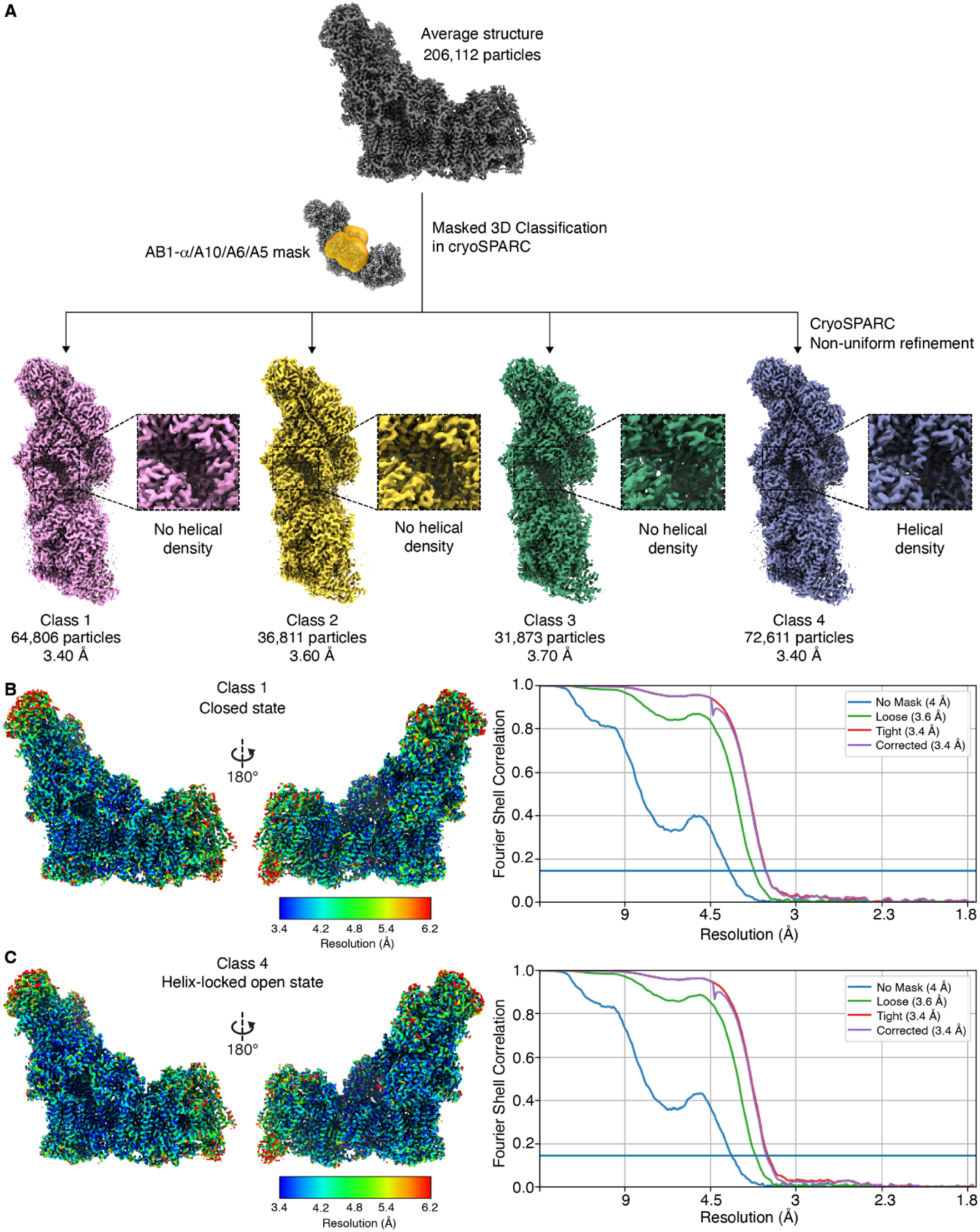
Identification of helix-locked open state. Masked classification of 206,112 *Dm*-CI particles using mask around the hinge region (NDUFA10, NDUFA5, NDUFA6 and NDUFAB1-α) at the PA/MA interface of *Dm*-CI resulted in four distinct classes of particles, two major classes (Class1 and Class 4) and two minor classes (Class 2 and Class 3). Major class 1 has no of density for the N-terminal region of NDUFS4 (shown in inset) (closed state) and major class 4 has clear density for the N-terminal region of NDUFS4 (locked open state). Minor classes 2 and 3 have no density for the N-terminal region of NDUFS4 (shown in inset). (B C) Local resolution maps and Fourier shell correlation (FSC) curves: closed state (B) and locked open state (C). Local resolution plotted on refined maps and FSC curves (gold standard FSC=0.143 for resolution estimation). Resolution scale bars are shown.

**Figure 4-figure supplement 2.**
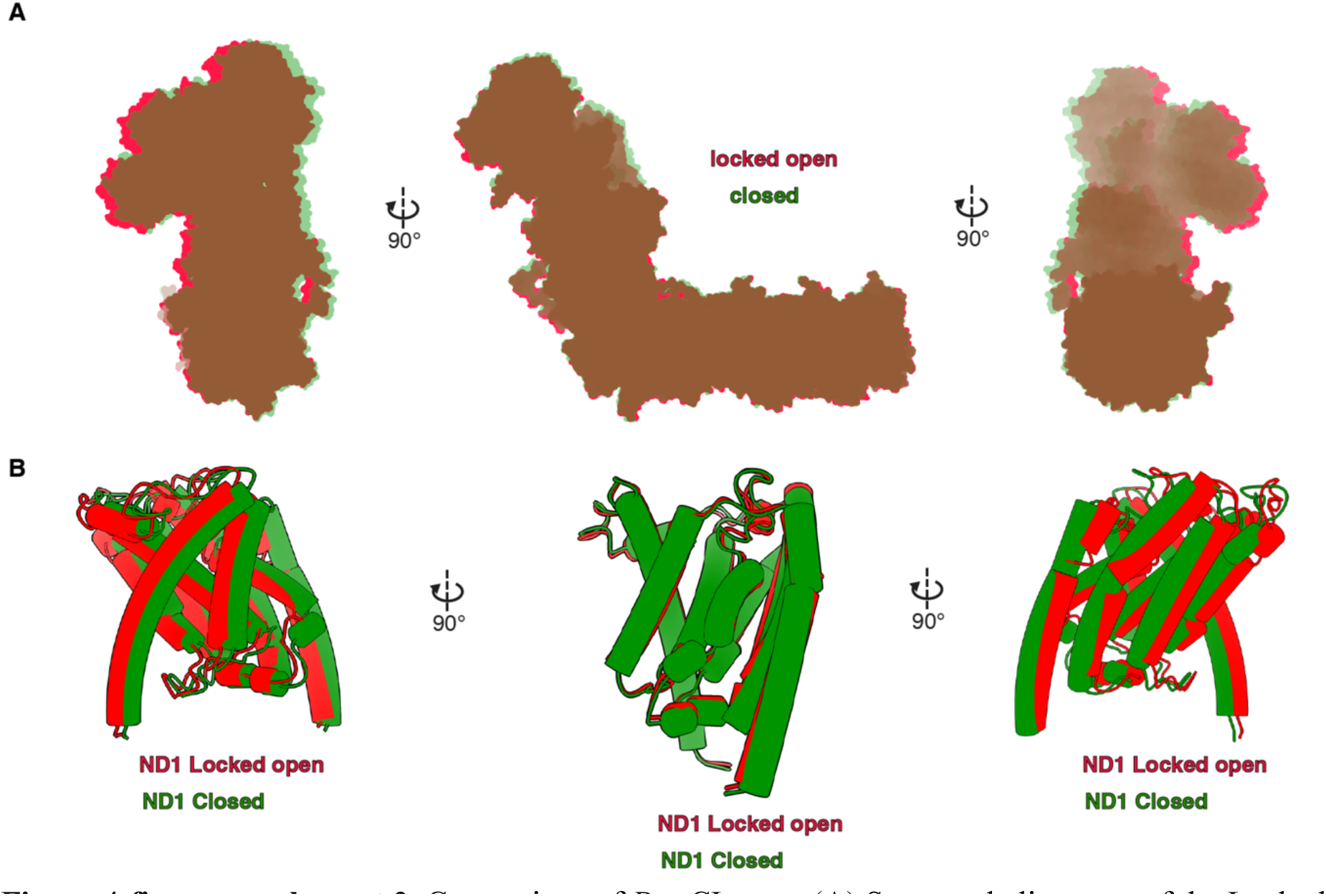
Comparison of *Dm*-CI states (A) Structural alignment of the Locked open state of Dm-CI (red) with the closed state of Dm-CI (green). The structures are aligned by ND5 subunit. (B) ND1 of Dm-CI in the locked open (red) and closed (green) state as seen in structures aligned by ND5 (above) are shown as cartoons.

**Figure 4-figure supplement 3.**
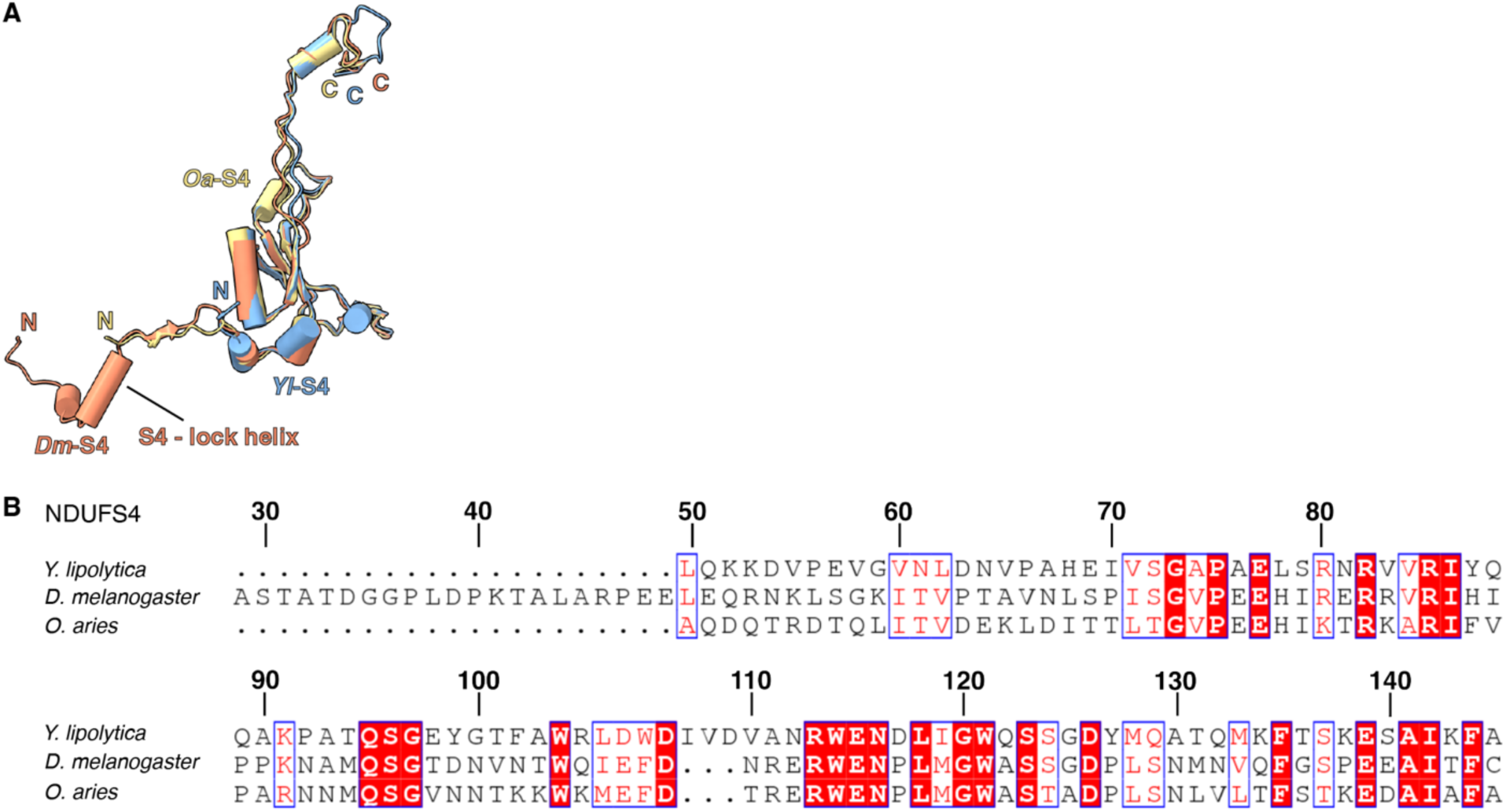
NDUFS4 comparison. (A) Structural alignment of N-terminal region NDUFS4 from *Y. lipolytica* (PDB: 6YJ4), *D. melanogaster* (this study) and *O. aries* (PDB: 6ZKC) with signal sequence removed is shown. (B) Sequence alignment of NDUFS4 from *Y. lipolytica*, *D. melanogaster* and *O. aries*. Numbered according to *Dm*-NDUFS4.

**Figure 4-figure supplement 4.**
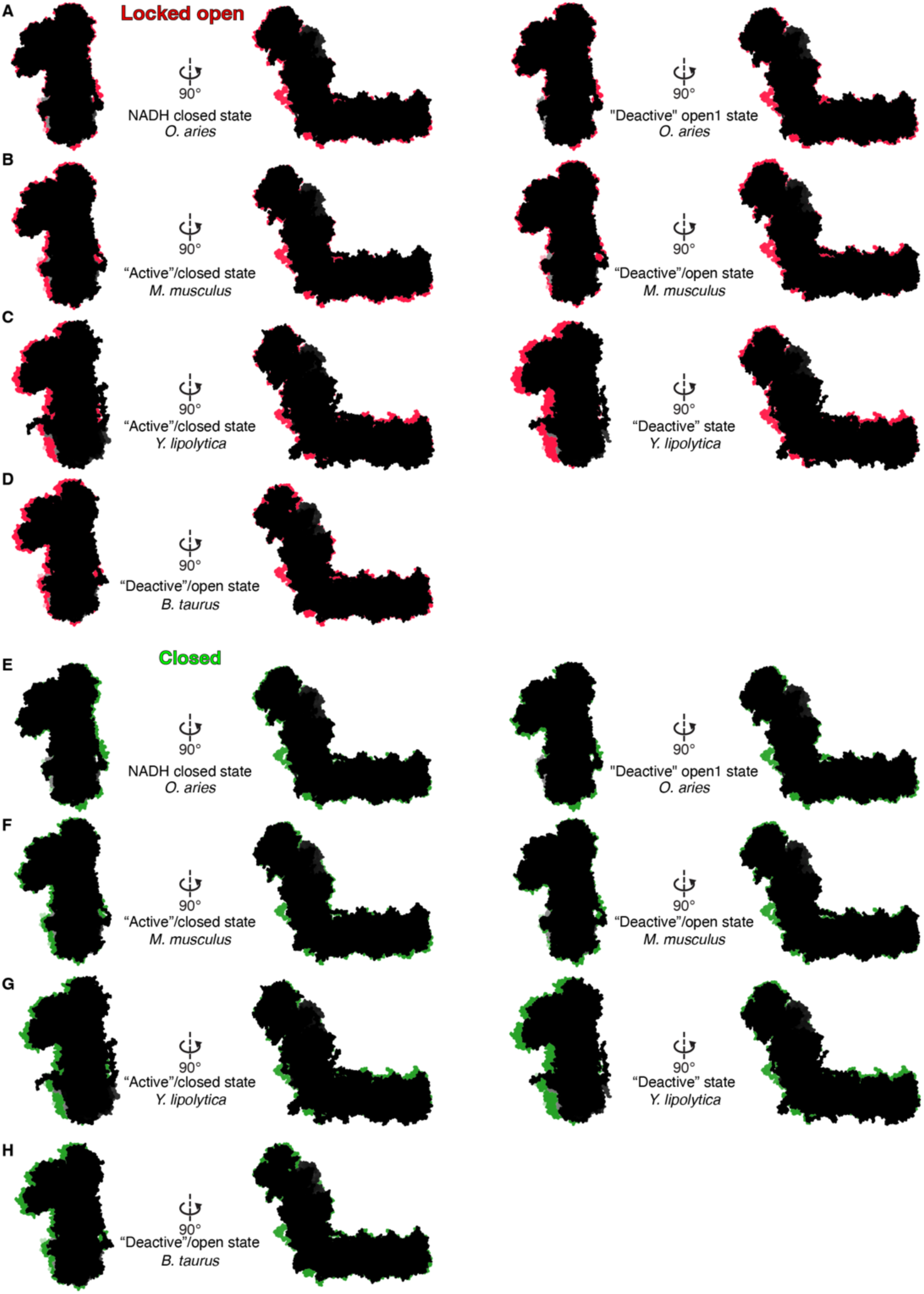
Comparison of helix-locked open state and closed state to states of other CI. (A-D) Structural alignment of Dm-CI locked open state structure (red) with the active and deactive states of CI from other organisms (black): NADH closed state of *O. aries* (PDB: 6ZKG) and deactive complex open1 of *O. aries* (PDB: 6ZKS) (A), active state of *M. musculus* (PDB: 6G2J) and deactive state of *M. musculus* (PDB: 6G72) (B), active state of *Y. lipolytica* (PDB: 6RFR) and deactive state of *Y. lipolytica* (PDB: 6GCS) (C), deactive state of *B. taurus* (PDB: 5O31) (D). CI structure is shown as surface. (**E-H**) Structural alignment of *Dm*-CI closed state structure (green) with the active and deactive states of CI from other organisms (black): NADH closed state of *O. aries* (PDB: 6ZKG) and deactive complex open1 of *O. aries* (PDB: 6ZKS) (E), active state of *M. musculus* (PDB: 6G2J) and deactive state of *M. musculus* (PDB: 6G72) (F), active state of *Y. lipolytica* (PDB: 6RFR) and deactive state of *Y. lipolytica* (PDB: 6GCS) (G), deactive state of *B. taurus* (PDB: 5O31) (H). CI structure is shown as surface. Accessory subunits are not shown for clarity.

**Figure 5-figure supplement 1.**
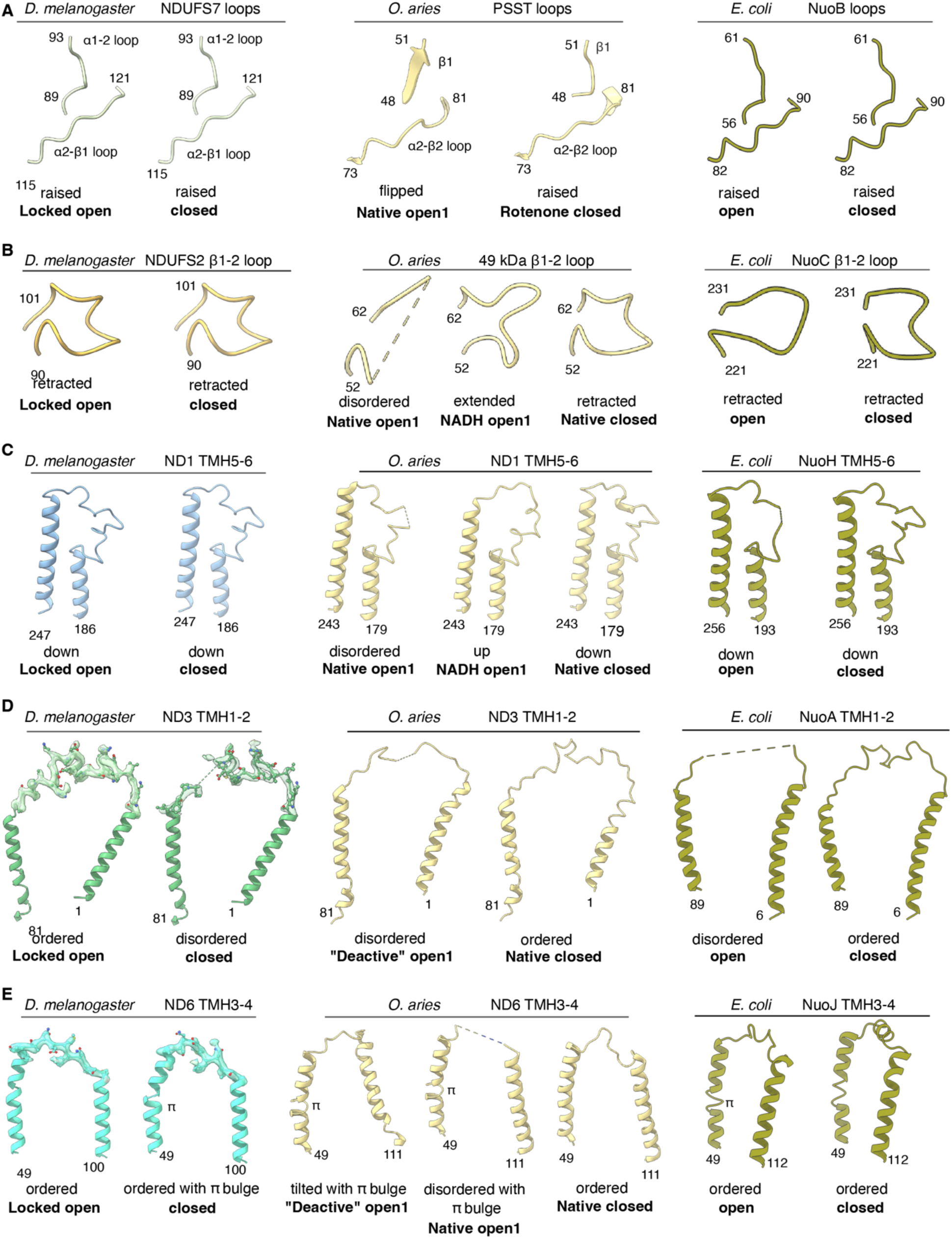
Structural comparison of the Q-site adjacent loops of *Dm*-CI in the Locked-open and closed states with the confirmations of *O. aries* and *E. coli*. (**A**) NDUFS7 loops of *D. melanogaster* in locked open and closed states (this study), flipped confirmation of PSST loops from *O. aries* CI (native open1, PDB: 6ZKP), raised confirmation of PSST loops from *O. aries* CI (rotenone closed, PDB: 6ZKK), *E. coli* (open, 7Z83) and (closed, 7Z80) states. (**B**) NDUFS2 β1-2 loop. NDFS2 β1-2 loop of *D. melanogaster* in locked open state and closed states (this study), disordered confirmation of *O. aries* CI (native open1, PDB: 6ZKP), extended confirmation of *O. aries* CI (NADH open1, PDB: 6ZKH), retracted confirmation of 49kDa subunit loop from *O. aries* CI (native closed, PDB: 6ZKO), *E. coli* (open, 7Z83) and (closed, 7Z80) states. (**C**) ND1 TMH5-6. *D. melanogaster* ND1 TMH5-6 loop in locked open and closed state, the disordered state of ND1 loop as of *O. aries* CI (native open1, PDB: 6ZKP), up confirmation of *O. aries* CI (NADH open1, PDB: 6ZKH), down confirmation of ND1 subunit of *O. aries* (native closed, PDB: 6ZKO), *E. coli* (open, 7Z83) and (closed, 7Z80) states. **(D)** ND3 TMH1-2 loop. ND3 TMH1-2 loop of *D. melanogaster* in locked open and closed states, disordered confirmation from *O. aries* (deactive open1, PDB:6ZKS), ordered confirmation from *O. aries* (native closed, PDB 6ZKO), E. *coli* (open, 7Z83) and (closed, 7Z80) states, **(E)** ND6 TMH3-4 loop. *D. melanogaster* ND6 TMH3-4 loop in locked open and closed states, the tilted configuration of CI from *O. aries* ND6 TMH4 with π bulge as seen in the open state of deactive CI from *O. aries* (deactive open 1, PDB: 6ZKS) disordered confirmation of CI with π bulge of *O. aries* (native open1, PDB: 6ZKP), ordered confirmation of *O. aries* (native closed, PDB: 6ZKO), *E. coli* (open, 7Z83) and (closed, 7Z80) states.

**Figure 5-figure supplement 2.**
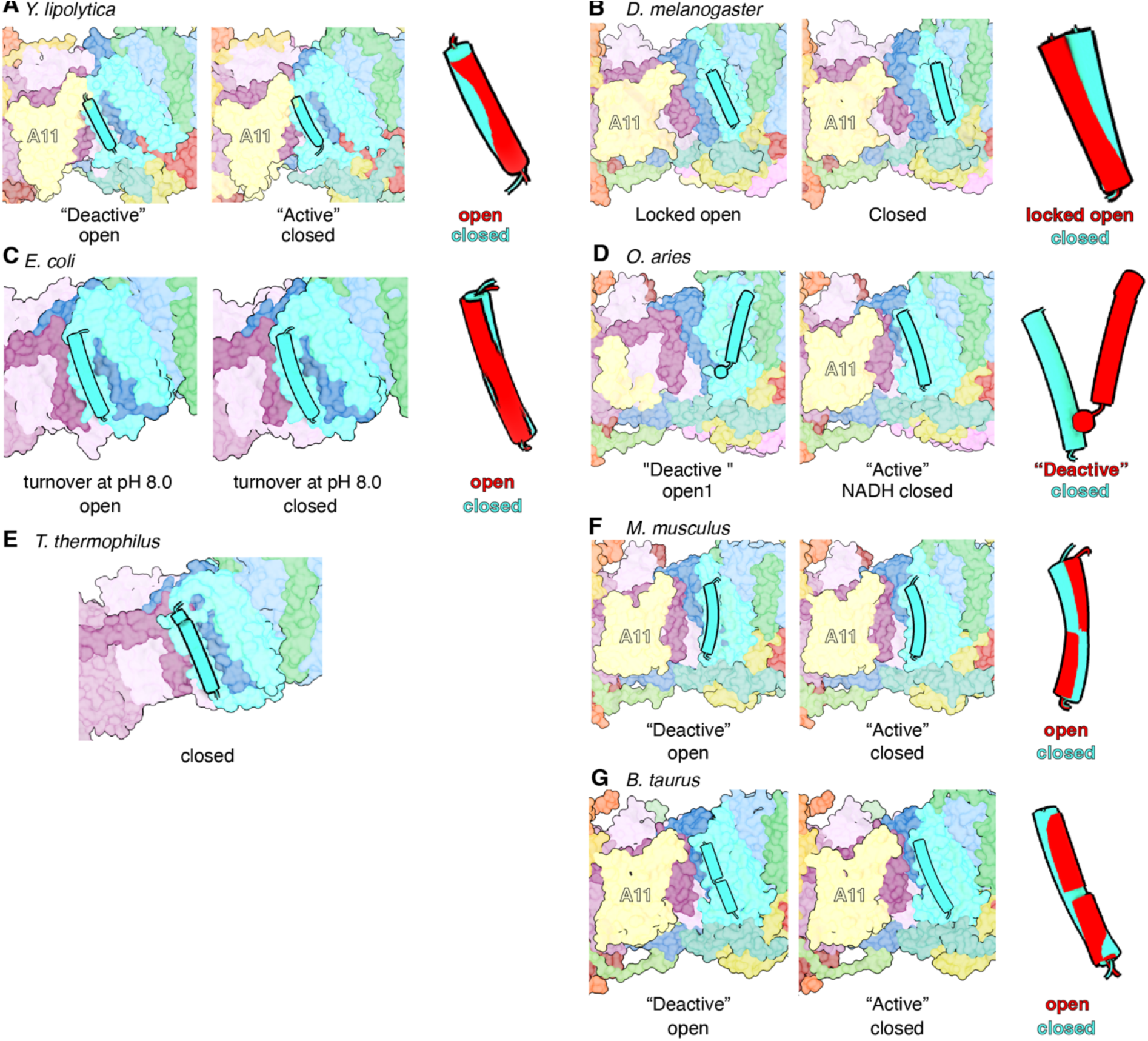
Comparison of TMH4^ND6^ orientation in the open and closed states across different organism**s.** (A) TMH4^ND6^ orientation in open (PDB:6GCS) and closed state (PDB:6RFR) of *Y. lipolytica*. TMH4^ND6^ relative orientation in structurally aligned ND6 of the two states is shown in the right panel. (B) TMH4^ND6^ orientation in locked open and closed state (this study) of *D. melanogaster*. ND6 TMH4 relative orientation in structurally aligned ND6 of the two states is shown in the right panel. (C) TMH4^ND6^ orientation in open (PDB:7Z83) and closed state (PDB:7Z80) of *E. coli*. TMH4^ND6^ relative orientation in structurally aligned ND6 of the two states is shown in the right panel. (D) TMH4^ND6^ orientation in open (PDB:6ZKS) and closed state (PDB:6ZKG) of *O. aries*. TMH4^ND6^ relative orientation in structurally aligned ND6 of the two states is shown in the right panel. (E) TMH4^ND6^ orientation in *T. thermophilus* (PDB:7Z83) is shown. TMH4^ND6^ is shown as a cartoon, the rest of CI is shown in colored surface. (F) TMH4^ND6^ orientation in open (PDB:6G72) and closed state (PDB:6G2J) of *M. musculus*. TMH4^ND6^ relative orientation in structurally aligned ND6 of the two states is shown in the right panel. (G) TMH4^ND6^ orientation in open (PDB:5O31) and closed state (PDB:5LC5) of *B. taurus*. TMH4^ND6^ relative orientation in structurally aligned ND6 of the two states is shown in the right panel.

**Figure 5-figure supplement 2.**
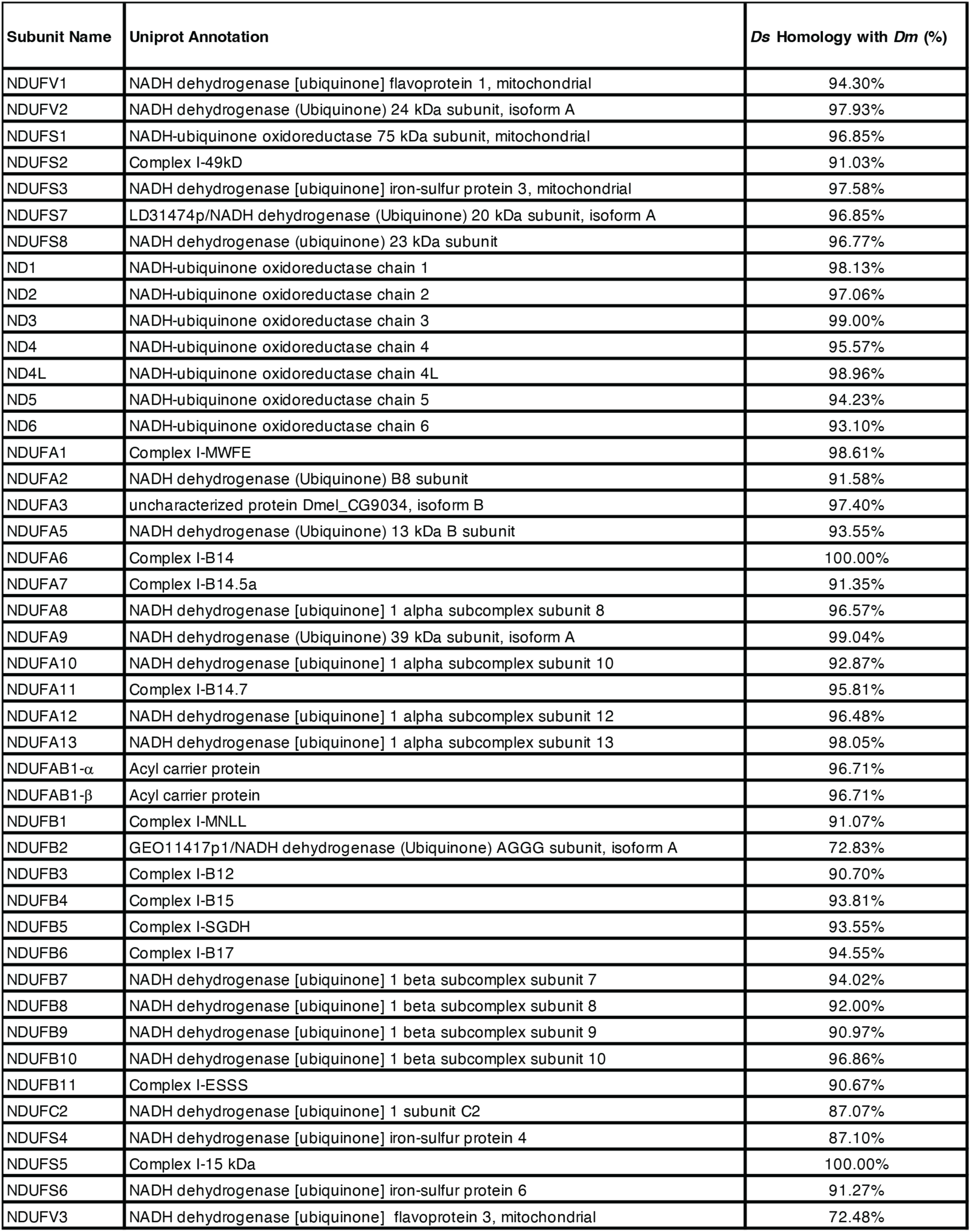
Sequence homology between CI subunits of *D. melanogaster* and *D. suzukii*.

